# Systematic genome-wide discovery of host factors governing bacteriophage infectivity

**DOI:** 10.1101/2024.04.20.590424

**Authors:** Chutikarn Chitboonthavisuk, Cody Martin, Phil Huss, Jason M. Peters, Karthik Anantharaman, Srivatsan Raman

**Affiliations:** Department of Biochemistry, University of Wisconsin-Madison, Madison, Wisconsin, USA; Department of Bacteriology, University of Wisconsin-Madison, Madison, Wisconsin, USA; Microbiology Doctoral Training Program, University of Wisconsin-Madison, Madison, Wisconsin, USA; Department of Chemical and Biological Engineering, University of Wisconsin-Madison, Madison, Wisconsin, USA; Pharmaceutical Sciences Division, School of Pharmacy, University of Wisconsin-Madison, Madison, Wisconsin, USA; Great Lakes Bioenergy Research Center, University of Wisconsin-Madison, Madison, Wisconsin, USA; Department of Medical Microbiology and Immunology, University of Wisconsin-Madison, Madison, Wisconsin, USA; Center for Genomic Science Innovation, University of Wisconsin-Madison, Madison, Wisconsin, USA; Department of Integrative Biology, University of Wisconsin-Madison, Madison, Wisconsin, USA

## Abstract

Bacterial host factors regulate the infection cycle of bacteriophages. Except for some well-studied host factors (e.g., receptors or restriction-modification systems), the contribution of the rest of the host genome on phage infection remains poorly understood. We developed ‘PHAGEPACK’, a pooled assay that systematically and comprehensively measures each host gene’s impact on phage fitness. PHAGEPACK combines CRISPR interference with phage packaging to link host perturbation to phage fitness during active infection. Using PHAGEPACK, we constructed a genome-wide map of genes impacting T7 phage fitness in permissive *E. coli*, revealing pathways previously unknown to affect phage packaging. When applied to the non-permissive *E. coli* O121, PHAGEPACK identified pathways leading to host resistance; their removal increased phage susceptibility up to a billion-fold. Bioinformatic analysis indicates phage genomes carry homologs or truncations of key host factors, potentially for fitness advantage. In summary, PHAGEPACK offers valuable insights into phage-host interactions, phage evolution, and bacterial resistance.

## Introduction

Bacteriophages (or “phages”), viruses that infect bacteria, rely on their bacterial hosts for replication and successful completion of their infection cycle. During the infection cycle, phages undergo a complex process involving adsorption, genome delivery, transcription of phage genes, assembly of new virions, and finally, the release of phage progeny. This infection cycle involves interactions with numerous host genes. Some host genes may facilitate phage infection by acting as receptors, supplying transcriptional and translational machinery, or enabling metabolic and nutrient uptake that phage appropriates. Other host genes may impede infection by producing antiphage components, restriction-modification enzymes, or mechanisms obstructing critical steps in the phage life cycle(1–3). The complex interactions between phages and hosts form the foundation of a dynamic evolutionary race that has unfolded over billions of years(3, 4). Despite decades of phage research, a comprehensive understanding of the contribution of each bacterial gene in the phage replication cycle remains poorly understood(4).

While hypothesis-driven research has successfully elucidated the roles of several host genes, the role of the vast majority of the host genome toward phage replication remains uncharacterized. Early screens on defining these relationships mainly relied on untargeted forward genetics, which is restricted to identifying genes with large effects on fitness (e.g. receptors), not a comprehensive global view. High-throughput targeted approaches, such as CRISPRi, have been employed to comprehensively phenotype genes relevant to phage infectivity(5–7). However, it is crucial to note that these studies have relied on host-based screens, where the impact of host genes on phages is measured through the changes in host fitness and survival. While an increase in host survival can indicate a decrease in phage fitness, this effect is still limited to host genes with a larger, pronounced effect on phage fitness, such as those involved in receptor recognition or specialized host defense mechanisms like restriction-modification and abortive infections(1, 2, 8–11). Capturing the effect of other host genes involved in phage reproduction, such as transcription, translation, codon usage preferences, and genome replication machinery, is challenging because these genes influence phage fitness without strongly affecting host viability(12–15). Ideally, perturbations to the host should be linked to phage fitness. However, linking host perturbation (via CRISPRi or transposon mutagenesis) to phage fitness is challenging because the host perishes after an active phage infection, and as a result, genome-wide screening approaches that characterize the relationship between host factors and phage replication have proven challenging to develop.

To address this challenge, we devised an approach to directly assess how disrupted host genes influence the ability of the phage to make phage progeny. We present ‘PHAGEPACK’ (**P**hage-**H**ost **A**nalysis using **Ge**nome-wide CRISPRi and phage **PACK**aging), a pooled assay for directing phage-based quantification and systematically characterizing host factors influencing phage fitness. PHAGEPACK establishes a direct link between host gene perturbation and phage activity by combining CRISPR interference (CRISPRi) with a phage packaging system. To prevent the loss of perturbation signal due to host lysis, we placed the sgRNA responsible for host gene disruption on a plasmid containing phage packaging sequence. The sgRNA is packaged into progeny phage particles and allows us to track changes in phage activity corresponding to each bacterial gene without signal loss during bacterial cell lysis (Figure 1A).

**Figure 1.**
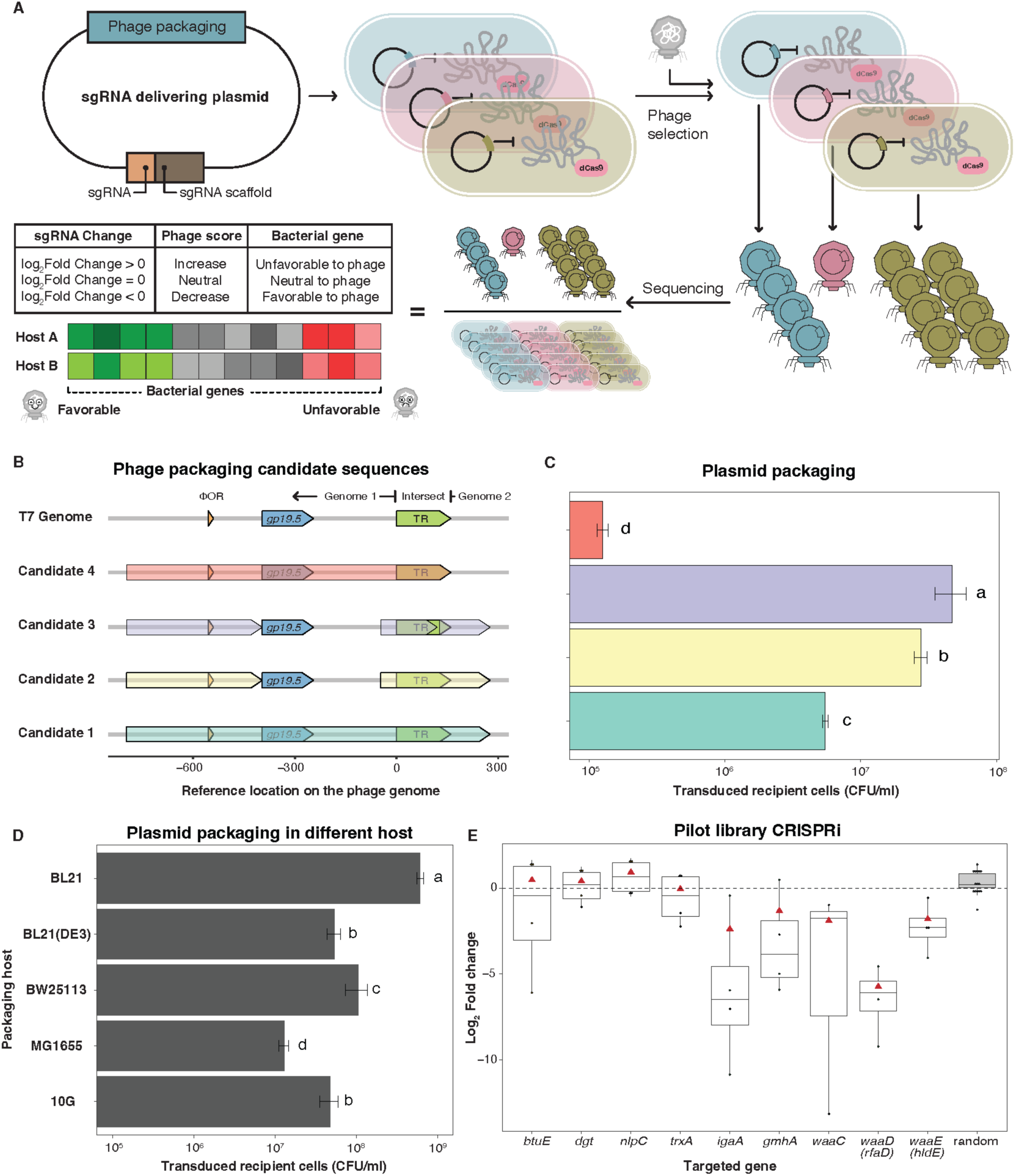
PHAGEPACK integrates CRISPRi and phage packaging strategies. (A) Schematic illustration of PHAGEPACK screening experiment. The sgRNA delivery plasmid, featuring a plasmid packaging sequence for phage packaging, is designed to target individual bacterial genes for a knockdown. The pooled experiment involves the simultaneous knockdown of bacterial genes using this plasmid with specific sgRNAs in *dCas9* integrated host. Phage selection is conducted, and the supernatant, containing plasmid-packaged phage undergoes sequencing. The relative abundance of each sgRNA in both phages and the bacterial population without phage selection is quantified. The sgRNA fold change (FC) is calculated and log-transformed, categorizing the role of bacterial genes on phage score as favorable (log_2_FC<0, green), neutral (log_2_FC=0, grey), or unfavorable (log2FC>0, red). (B) Four phage packaging sequences with varying coverage on the T7 genome were evaluated to test packaging abundance. Each color represents each candidate. (C) Quantification of plasmid packaging abundance across candidates is determined by measuring the absolute abundance of packaged plasmids through transduction in a *ΔtrxA* recipient, represented in CFU/ml. (D) Chosen candidate was evaluated for packaging abundance in various *E. coli* strains (K12 (BW25113 and MG1655),B (BL21(DE3) and BL21), and 10G. Difference of plasmid packaging abundance was determined by Tukey’s HSD test at the *p-*value of 0.05 in C and D. (E) A pilot library containing selected sgRNAs targeting bacterial genes, including *btuE*, *dgt*, *nlpC, trxA*, *igaA*, *gmhA*, *waaC*, *waaD* (*rfaD*), and *waaE* (*hldE*), and non-targeted or randomized sgRNA (grey). Boxplot shows the phage score represented in log_2_FC with the log-transformed of mean FC (▴).

We applied PHAGEPACK to create a genome-wide quantitative map of the impact of *E. coli* genes on T7 phage fitness. Besides validating known T7-*E. coli* interactions, this screen revealed dozens of potential host factors that had both facilitated and impeded phage replication that, prior to this study, were not associated with T7 fitness. These included genes involved in iron-sulfur cluster biogenesis, chemical modification of tRNA, codon usage preferences, post-transcriptional regulation, and toxin-antitoxin systems. In addition to the permissive host, PHAGEPACK is uniquely suited for evaluating phage-host relationships in non-permissive hosts, as these hosts typically produce minimal or no phage progeny, making even minor increases in phage fitness detectable. PHAGEPACK differs from host-based screens because it directly quantifies the increase in phage fitness, rather than relying on the loss of signal from a disrupted host. We applied PHAGEPACK to a foodborne pathogen, *E. coli* O121, that is not permissive to T7 infection, with the goal of uncovering host factors that impeded T7 infectivity. PHAGEPACK revealed that pathways involved in bacterial outer layer composition and diverse internal pathways associated with core functions are key major barriers to T7 infection in *E. coli* O121. Knocking down these genes increased the infectivity of T7 by up to a billion-fold.

Finally, the results of PHAGEPACK also allowed us to explore and understand viral evolution in relation to these host factors. Bioinformatic analysis revealed that viral genomes are enriched with homologs of host genes that increase phage fitness as well as truncated or mutated versions of genes that decrease phage fitness. These results support a dominant negative hypothesis where phages may express malformed proteins that interfere with host genes that would otherwise reduce phage fitness. In summary, PHAGEPACK is a powerful tool for understanding the relationship between phages and bacterial hosts, offering valuable insights into host factors influencing phage dynamics and affecting phage fitness.

## Results

### PHAGEPACK links host perturbations to phage fitness

Phage-induced lysis of the host cell is a significant challenge in creating a pooled screen that connects phage fitness to the host because host markers (e.g., barcodes or guide RNA) will be lost. To overcome this challenge, we exploited the natural phage packaging system. A phage packaging sequence is a specific DNA sequence, often located at the ends of a phage genome, serving as a recognition signal for the phage packaging machinery to cleave the phage genome into a unit-length form and package it into the capsid(16).

Generally, most dsDNA phages replicate their genome as concatemers, which are recognized, cleaved, and packed into phage capsids by specific regions. While these regions can vary among phages – such as *cos*, *pac*, or direct terminal repeat (DTR) sequences – they are commonly located at the DNA termini(16). Packaging sequences and schemes have been characterized in various phages. Therefore, PHAGEPACK can be generalized to other phages by leveraging the packaging system.

T7 phage uses a 160-bp terminal repeat (TR) for packaging, which is a redundant part of the 5’-and 3’-end of the genome concatemerized during replication(17). We incorporated the single-guided RNA (sgRNA) library into a plasmid containing the T7 packaging sequence (TR and 3’-end origin of replication). When T7 infects a host cell carrying a specific sgRNA, that sgRNA is packaged into an empty phage. The impact of a gene knockdown can be quantitatively assessed by counting the number of phages containing that sgRNA. For example, knocking down an essential gene for phage replication will result in fewer phages containing the corresponding sgRNA, while knocking down a gene detrimental to phage replication will lead to a higher number of phages containing the corresponding sgRNA. A functional score is then assigned to each bacterial gene based on the relative abundance of each sgRNA to quantify its influence on phage infection.

We overcame several challenges in developing PHAGEPACK (Figure 1). First, we confirmed that introducing a packaging sequence would indeed produce phages containing the plasmid. Second, we identified the specific packaging sequence that yielded the highest abundance of packaged phages to enhance the signal-over-noise. Third, we established generalizability by demonstrating the chosen packaging sequence was effective on different *E. coli* strains. Finally, we validated that quantifying phage with sgRNA accurately reflected the underlying biological role of host factors in phage infectivity.

DNA can be packaged into phages by using a packaging sequence containing the origin of replication (ΦOR) on the T7 genome and the concatemeric junction within each TR(17–19). We selected four packaging sequences with different concatemeric junctions and their surrounding regions to assess packaging (Figure 1B). We initially validated the packaging of all four packaging sequences in permissive *E. coli* 10G. The abundance of packaged phages was quantified through transduction in an *E. coli* Δ*trxA* mutant(20). Of the four constructs, we found that candidate 3 (pT7cand03) produced the greatest number of transduced cells, reflecting the number of packaged phages (Figure 1C, Table S3). To assess the generalizability of pT7cand03 on different *E. coli* variants, we tested the packaging of pT7cand03 on *E. coli* K12, MG1655 and BW25113, B strains, BL21(DE3) and BL21(REL606), and 10G. We found that all five strains can support packaging of pT7cand03 following wildtype T7 infection, with comparable abundances of packaged phages (Figure 1D).

Next, we devised a pilot screen to assess whether quantifying sgRNA packaged within phage accurately reflects the impact of a host gene on phage fitness. We created an *E. coli* strain carrying genomic *dCas9* and confirmed the knockdown efficiency of four genes – three genes involved in T7 infection *lpcA*, *trxA*, and *waaC* and an *rfp* control (Supplementary Figure S1A and S1B)(21). For the pilot screen, we chose nine bacterial genes, including six genes known to impact phage fitness (*igaA*, *gmhA, waaC, waaD* (*rfaD*), *waaE* (*hldE*), and *trxA*) and three genes that have no impact (*btuE*, *dgt* and *nlpC*) (Figure 1E, Table S3). The former group included genes essential to T7 such as *igaA*, a regulator of the envelope stress response Rcs system, and *gmhA, waaC, waaD* and *waaE*, required for lipopolysaccharide (LPS) biosynthesis(5). The *trxA* gene is used for T7 DNA polymerase processivity and is essential for plaque production but has less effect on fitness in liquid culture(22). In total, this pilot experiment used 51 sgRNAs (9 genes with 4 sgRNAs per gene) and 15 randomized sgRNAs as a control.

To determine a functional score for each host gene on phage fitness, we computed the fold change ratio (FC) of the abundance of sgRNA in phage to their abundance in the bacterial host population after deep sequencing, using the standard edgeR calculations as employed in other CRISPR genetic screens(23). The sgRNA causing a drastic loss or a depletion of bacterial fitness is filtered out during the edgeR process. In the pooled PHAGEPACK screen, *igaA, gmhA, waaC, waaD* and *waaE* exhibited strong negative log_2_FC scores, validating their known roles in promoting phage infection (Figure 1E). The log_2_FC scores for *trxA* were comparable to randomized controls, supporting previous results that indicate this gene is less essential in early stages of infection in liquid culture. Log2FC scores of *btuE, dgt* and *nlpC* were comparable to randomized sgRNAs, further supporting their lack of a known role in phage infection. In summary, these results showed that PHAGEPACK can quantitatively recapitulate known host factor-phage relationships and can be applied to characterize host-phage interactions genome-wide.

### PHAGEPACK reveals genome-wide phage fitness scores

We scaled up PHAGEPACK genome-wide by designing a sgRNA library containing 18,471 targeted sgRNAs (four sgRNAs per gene) for BW25113 (K12-strain) and BL21 (B-strain) in addition to 1,000 randomized sgRNAs as control (Table S2). The *dCas9* was placed under inducible control to prevent the depletion of sgRNAs targeting genes with a strong impact on bacterial fitness before phage selection. After transforming the library into both strains, we observed a very high library coverage (>99.9%), with only a small fraction of genes missing – approximately 8 genes in BW25113 and 15 genes in BL21. Cells containing the pre-selection library were split into two populations. The first population, induced to express *dCas9* (without phage application), provided a reference by accounting for changes in host fitness. The second *dCas9*-induced population was infected with T7 at an MOI of 0.01 to minimize cross-infectivity by multiple phages. After 30 minutes of incubation, representing approximately one T7 replication cycle, we sequenced the sgRNA pool from the progeny phage and the reference host cells not subjected to T7 infection (Supplementary S2B and S2C). As described earlier, we computed the FC ratio of sgRNA abundance packaged in phages to their abundance in the bacterial host population using edgeR, averaging over all sgRNAs for a gene.

We conducted several preliminary checks before analyzing the full dataset. First, we verified that the results from this comprehensive genome-wide screen were consistent with our pilot screen (Figure 2A). The genes *gmhA*, *waaD* (*rfaD*), and *waaE* (*hldE*) significantly reduced phage fitness, while *waaC* and *igaA* also reduced phage fitness but with less significance when compared to the larger pool of a thousand randomized sgRNAs. Gene *trxA* consistently showed a minor impact on phage fitness. Genes *btuE*, *dgt*, and *nlpC* did not impact phage fitness. The verification was consistent in *E. coli* BL21, with genes *gmhA*, *waaC*, *rfaD*, and *hldE* significantly reducing phage fitness and *trxA* showing a lesser impact on phage in the liquid culture. Gene *igaA* did not have a significant impact on the phage in *E. coli* BL21. Overall, the effect of gene knockdowns was consistent with the results of the pilot study. The randomized sgRNA library showed no impact on phage fitness (Supplementary Figure S3), and the distribution of scores for the randomized sgRNAs was used as a null distribution for evaluating the statistical significance of targeted genes. The full dataset revealed large sets of genes that influence phage activity in both *E. coli* K12 and B strains (Figure 2B and 2C).

**Figure 2.**
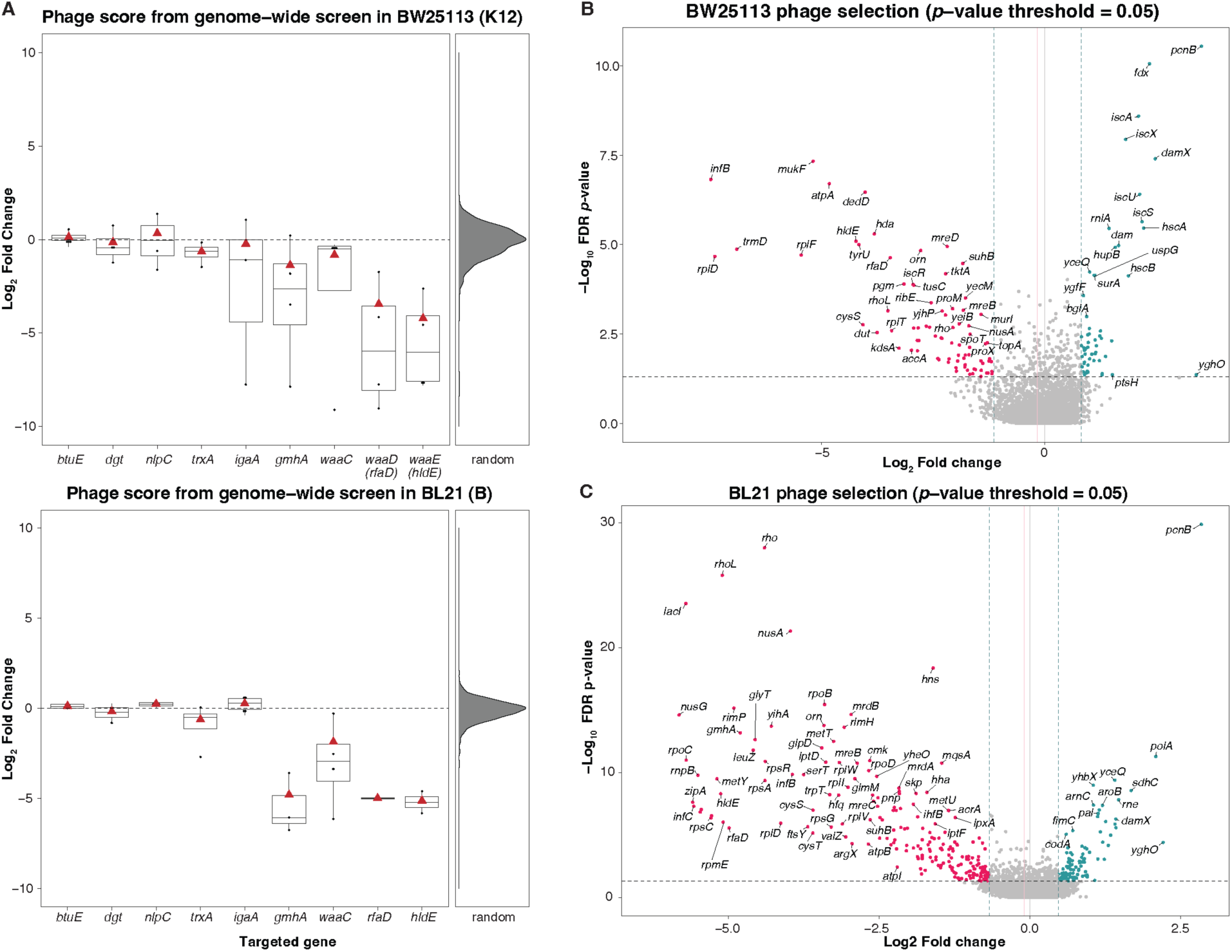
Genome-wide impact of host genes on T7 fitness. (A) Boxplots showing the phage score (represented in log_2_FC) and log-transformed of mean FC (▴) resulting from CRISPRi knockdown of each gene, as included in the pilot experiment for both *E. coli* K12 (top) and B strains (bottom). The right panel illustrates the distribution of 1,000 randomly generated sgRNAs from the library. (B) Volcano plots illustrate score for each gene, with increased scores in green, decreased in red, and neutral in grey, resulting from a gene knockdown in *E. coli* K12. Genes were considered significant at a *p-*value<0.05 (horizontal dashed line; adjusted *p-*value by false discovery rate (FDR)). The vertical dashed line delineates the standard deviation of the randomized sgRNA distribution, while the solid line represents the mean of the randomized sgRNA distribution (pink), coinciding with a log_2_FC of 0 (grey). (C) The volcano plot illustrates the effect on phage score resulting from a gene knockdown in *E. coli* B.

PHAGEPACK identified many genes that significantly impact T7 fitness that have been previously identified in host-based screens(6, 21). These include host genes involved in LPS synthesis, such as *hldE*, *rfaD*, and *gmhA*, as well as other genes that influence phage activity, such as *tktA*, *iscR, waaJ*, and *pgm*(6).

PHAGEPACK revealed five host pathways as major players in governing T7 infectivity on *E. coli* through previously unknown mechanisms (Figure 3). These include iron-sulfur cluster biogenesis, codon usage, polyadenylation, cell morphology, and toxin-antitoxin system (TA). Genes involved in these mechanisms had the largest FC score with high statistical significance, both positively and negatively influencing phage fitness (Table S4).

**Figure 3.**
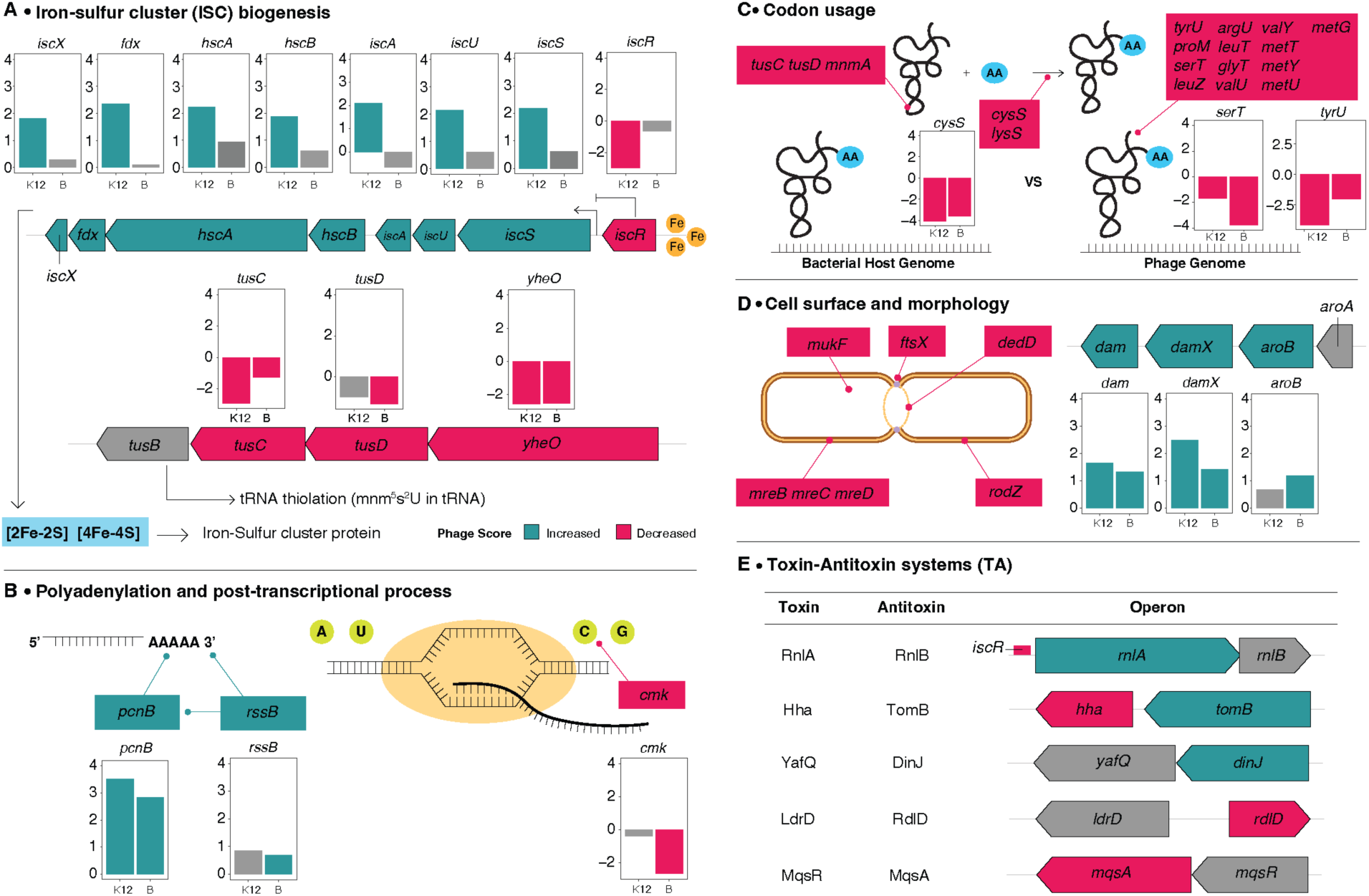
Novel host mechanisms influencing T7 phage infectivity. CRISPRi library experiment, *E. coli* K12 and B strains, reveals five potential mechanisms impacting T7 infection, including (A) iron-sulfur cluster (ISC) biogenesis, (B) polyadenylation and post-transcriptional process, (C) codon usage, (D) cell surface and morphology, and (E) toxin-antitoxin systems. Genes predicted to be involved in phage infection are categorized as beneficial (red), detrimental (green), or neutral (grey) to phage based on the knockdown result CRISPRi screen. Each individual bar plot represents the log_2_FC on the y-axis and *E. coli* strain on the x-axis.

#### Iron-sulfur cluster (ISC) biogenesis

The iron-sulfur cluster (ISC) operon consists of *iscRSUA*, *hscBA*, *fdx*, and *iscX* where *iscR* is the principal regulator of the operon(24). We found that knocking down each enzyme complex of the pathway, *iscSUA*, *hscBA*, *fdx*, and *iscX,* reduced phage fitness. Conversely, we found that knocking down the regulator of this operon, *iscR*, increased phage fitness (Figure 3A). This ISC housekeeping pathway, along with SUF and NIF pathways, plays a crucial role in maintaining Fe-S proteins, which account for approximately 3% of the *E. coli* proteome. Fe-S proteins have diverse roles, and are crucial for electron transfer in respiration, tRNA and rRNA modification, and transcriptional regulation(25, 26). Additionally, the disruption of individual genes leads to distinctive changes in the proteomic profile of these Fe-S proteins(27). ISC genes have been implicated in phage fitness previously in λ phage. ISC genes were found to affect the extent of thiolation of lysine-tRNA in *E. coli*, resulting in programmed ribosomal frameshifting to favor the translation of one λ protein over another, ultimately conferring a fitness advantage to λ phage(28). Our observation of decreased phage fitness upon knockdowns of genes *tusC, tusD*, and *mnmA*(29) suggests a potential impact of thiolation on T7 phage as well. Other possible mechanisms involve the regulatory role of *iscR*(28, 30). In T4 infection, it was demonstrated that the host IscR-regulated RnlAB TA system induces degradation of most mRNA during the late stage of infection. Consequently, T4 employs a Dmd inhibitor to counteract this host defense TA system(30, 31). Additionally, IscR has been shown to control biofilm production in response to iron-sulfur homeostasis, with more biofilm being produced in Δ*iscR* hosts(32). The indirect effector pathways of ISC and IscR, TA defense systems, and biofilm production may all influence the activity of T7 phage.

#### Polyadenylation and post-transcriptional process

The *pcnB* gene knockdown significantly enhanced phage fitness (Figure 3B). The *pcnB* gene encodes a poly(A) polymerase, an enzyme that polyadenylates transcripts in *E. coli*. The short poly(A) tails in *E. coli*, typically 10-40 bp, reduce mRNA half-life by promoting degradation through 3’ exonuclease activity(33, 34). Prior studies have shown that phage transcripts, T7 and λ are polyadenylated during the infection cycle(35–37). We hypothesize that knocking down *pcnB* likely increases the half-life of T7 transcripts, resulting in more phage progeny and higher phage fitness.

#### Codon usage

We identified a significant number of tRNA genes, such as *argU*, *glyT*, *valU*, *tyrU*, *proM*, *serT*, *metT*, *leuT* and tRNA ligase, *cysS* and *lysS*, that reduced phage fitness when knocked down (Figure 3C). This suggests that codons associated with these tRNAs may confer a fitness advantage to the phage potentially through enhanced translation efficiency of phage proteins or better adaptation to the endogenous environment of the host(13, 14, 38).

#### Cell surface and morphology

We found that knocking down genes involved cell division, cell wall synthesis, and the maintenance of cell shape and structure in *E. coli* – *dedD*, *ftsX*, *mreB*, *mreC*, *mreD* – reduced phage fitness (Figure 3D). Since these genes contribute to structural integrity, their knockdown may lead to abnormalities in the cytoskeleton and divisome complex and overall impairments in cell morphology. Changes in cell shape and surface structures may affect the availability or accessibility of host receptors, influencing the ability of phages to recognize and infect cells. Conversely, phage fitness increased with the knockdown of *damX*, a cell division protein located in the inner membrane. Increased phage fitness under *damX* knockdown could be due to the reduction of membrane integrity in the host(39–43), supporting the release of phage progeny. The strategy of phages disrupting the cell morphology to benefit their infection is common, for example, T7 Gp0.4 and λ Kil protein were shown to prevent FtsZ in *E. coli*, leading to a cell elongation(44, 45). Moreover, such changes could confer a competitive advantage to phage by releasing the infected cell from genome replication, thereby making more resources available for the phage.

#### Toxin-Antitoxin systems (TA)

The infectivity of T7 is significantly impacted by knocking down several TA systems, including *rnlA*, *hha/tomB*, *dinJ*, *rdlD*, and *mqsA* (Figure 3E). Host TA systems are stress response mechanisms to counteract phage predation through different mechanisms, such as triggering abortive infection, forming persister cells, inducing dormancy, and other adaptations. Specifically for RnlA, an mRNA endoribonuclease, we reasoned that its knockdown could lead to decreased phage mRNA degradation, supporting the production of phage progeny(30). Although our approach identified a significant impact of several TA genes, the lack of a detailed understanding of the mechanisms underlying TA systems limits nuanced speculation.

In summary, PHAGEPACK offers a systematic framework for discovering interactions between phage fitness and host gene activity. It unveils novel host pathways and mechanisms that influence phage fitness and is a powerful tool for hypothesis generation. Employing phage-based quantification instead of host-based quantification enables the discovery of host factors that exert both large and small positive or negative effects on phage activity, revealing new host genes and pathways relevant to phage fitness.

### PHAGEPACK reveals factors conferring phage resistance in a non-permissive strain

We next sought to determine if PHAGEPACK could be used to identify factors that impede phage infection in non-permissive strains. A significant hurdle in the broader application of phages for antibacterial therapy lies in their restricted host range. Phages target specific receptors on bacteria to initiate infection, while bacteria, in turn, develop countermeasures to thwart such infections. For example, T7 phage binds to rough bacterial LPS to infect certain *E. coli* strains. However, T7 does not interact with smooth LPS, and cannot infect *E. coli* strains possessing full-length O-antigen(46). Similarly, the infection of *Klebsiella* phages can be challenged depending on bacterial outer structures as K-and O-antigens(47). Minor genomic variations between strains can create formidable barriers to phage infectivity, manifested either through the host acquiring anti-phage systems or alterations in the host’s receptor profile(9, 20, 48).

We employed PHAGEPACK to identify host factors contributing to T7 phage resistance to non-permissive foodborne pathogen *E. coli* O121. This clinical strain can cause intestinal bleeding and hemolytic-uremic syndrome as one of the “Big Six” non-O157 serotypes associated with worldwide food-borne infections(49). We expected PHAGEPACK to be uniquely suited to evaluating non-permissive hosts, as any gene knockdown that produces phage progeny would generate a high signal and can thus be easily identified.

T7 phage does not infect *E. coli* O121 in tested conditions and has no perceivable plaque formation or effect on cell growth (Figure 4A, Table S5). To discover the host factors contributing to this resistance, we designed a sgRNA library comprising 18,583 targeted sgRNAs (four sgRNAs per gene) covering the genome of O121 and 1,000 randomized sgRNAs for controls (Table S2). As expected, most genes (98.8% of total genes) had a negative log_2_FC score, suggesting that knocking down these genes provides little or no fitness advantage to the phage compared to the randomized control (Supplementary Figure S7). The remaining genes gave moderate-to-high log_2_FC scores. We observed a larger error in moderately impact genes, compared to one whose knockdown significantly increased phage fitness, due to a lower abundance of sgRNAs detected after a phage selection (Supplementary Figures S8 and S10). We identified 28 genes with small but significant effect phage fitness with log_2_FC between −2.5 and 2 (Figure 4B and Supplementary Figure S9) and approximately 40 gene knockdowns resulted in a sharp increase in phage fitness with a log_2_FC greater than 2 (Figure 4B and Supplementary Figure S9).

**Figure 4.**
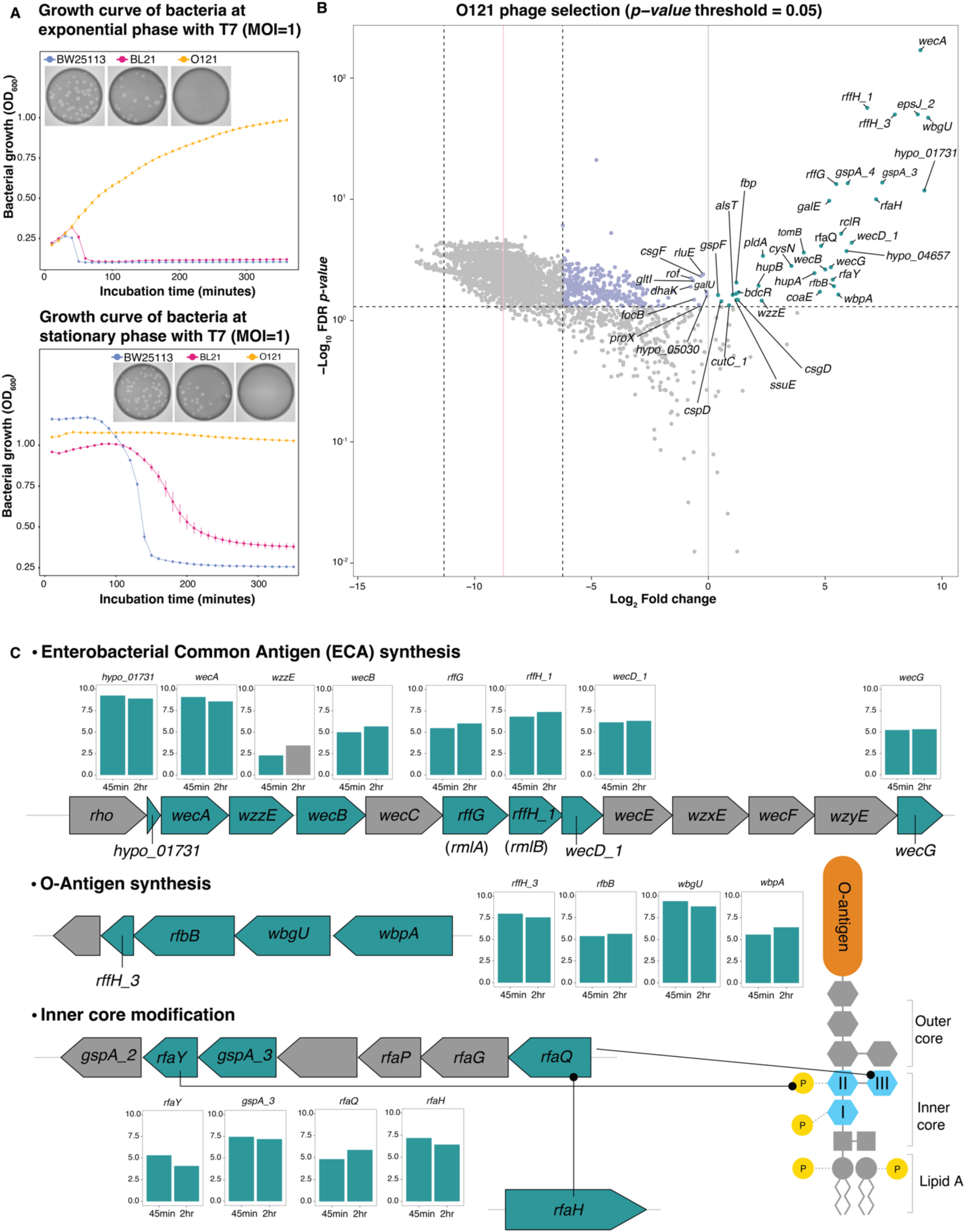
Mechanisms of T7 phage resistance in *E. coli* O121. (A) A bacterial growth curve (measured by OD_600_) illustrates the growth of *E. coli* BW25113 (blue), BL21 (red), and O121 (yellow) – under phage infection at the exponential phase (top) and stationary phase (bottom) at an approximate MOI of 1. (B) A volcano plot showing phage scores obtained after selection on *E. coli* O121, with significantly increased scores in green (log_2_FC>0), detectable but negative scores in purple (log_2_FC larger than the standard deviation of randomized dataset), and non-significant scores in grey. The horizontal dashed line indicates a confidence level at a *p-* value of 0.05, and four vertical lines represent mean of randomized sgRNAs (solid pink), its standard deviation (dashed), and zero (solid grey). (C) Bacterial mechanisms predicted to primarily influence T7 phage infections. Bacterial factors were categorized as unfavorable (green) or neutral (grey) to phage activity. Bar plots show phage scores observed in CRISPRi screen from both 45-minute and 2-hour selections. Three predicted mechanisms included Enterobacterial Common Antigen synthesis (ECA), O-antigen, and core LPS modification. Genes were depicted within their respective operons. The accompanying structure illustrates LPS components, including lipid A, inner and outer core LPS, and the O-antigen. Heptose I, II, and III were represented in I, II, and III labelled hexagons. P represents phosphate.

The strongest hits represent genes involved in shaping the outer surface structure of *E. coli* O121, including biosynthetic pathways of the Enterobacterial Common Antigen (ECA; *wecABDG*, *rffH* [*rmlA*], *rffG* [*rmlB*], *wzzE*), the O-antigen (*wecA*, *wbpA*, *wbgU*), and the core LPS (*rfaY* [*waaY*], *rfaQ* [*waaQ*], *rfaH*) (Figure 4C). Results for the core LPS contrast those on permissive strains. Knockdown of these genes may substantially affect the morphology of LPS and O-antigen in a cascading manner, increasing phage activity. Alternatively, since *rfaY, rfaQ,* and *rfaH* are involved in the phosphorylation and transfer of heptose to the core LPS structure, the loss of phosphorylation on heptose II and potential changes in membrane permeability could enhance phage infection(50). Previous studies showed that the disruption of these genes individually results in the loss of phosphorylation, thereby altering the negative charge of the membrane(51). Although this phosphorylation has no discernible impact on phages in permissive strains, even a slight loss or change in charge around the core LPS might enhance the accessibility of the phage to the LPS receptor. These results strongly suggest that surface modifications of *E. coli* O121 are the primary barrier to T7 activity. Knocking down genes involved in the biosynthesis of surface sugars significantly enhances T7 infectivity by likely disrupting or dismantling barriers to receptor recognition or orientation of the phage on the bacterial surface for successful infection.

Genes with moderate impacts on T7 infection encompass a diverse array of functions, extending beyond outer structure to potentially influence various stages of the phage lifecycle, including transcription, translation, and lysis processes. The genes related to transport (*ehaG, alsT*, *proX, gltI*), surface and cell structure (*dacC, gmm, minC*), and motility (*fliH, csgF*) are likely to be involved in the initial interaction of phage and the bacterial cell at the outer structure. While these host factors may not directly constitute the LPS receptor for T7, their presence on the bacterial cell membrane could potentially render them recognizable by T7, thus facilitating the infection(52, 53). Additionally, there are several genes that may be involved during phage replication. For instance, the knockdown of the transcriptional regulatory RclR(54), known as a redox-sensitive activator, could alter the host’s sensitivity and impact phage replication or lysis. Despite observing a significant increase in phage fitness, as evidenced by clear plaques, the underlying mechanism remains unclear. The genes *rluE* and *rihA* can impact phage transcription as they are involved in pseudouridine synthesis and the hydrolysis of uridine/cytidine to uracil/cytosine, respectively. Additionally, changes in metabolism may also affect phage activity, as potentially seen with DhaK, a dihydroxyacetone kinase pivotal in glycerol metabolism.

These findings offer crucial insights into the mechanisms of phage resistance within a non-permissive bacterial pathogen. Screening other pathogens using PHAGEPACK will generate a comprehensive catalog of potential resistance mechanisms, providing valuable guidance for phage therapy.

### Clonal validation of *E. coli* O121 host factors on T7 infectivity

To assess the impact of key *E. coli* O121 genes found by PHAGEPACK, we individually knocked down gene hits and validated the resulting changes in phage activity by plaque assay (Figure 5A, Table S6). Genes associated with ECA biosynthesis (*wecA*, *wecD*, *rffH*, and *wzzE*), the O-antigen (*wbpA* and *wbgU*), and core LPS modification (*rfaH* and *rfaY*) showed the most significant improvements in phage activity and resulted in nearly complete rescue of T7 activity, comparable to infection in permissive hosts. The largest gain observed was approximately a billion folds considered as a full rescue. Plaques formed on these hosts were clear, similar to the plaque formed on *E. coli* K12, suggesting unencumbered infection. Other genes, such as *alsT*, *minC*, *hypA*, *cysN*, and *hupA*, exhibited an approximate 4-7 log increase in phage activity while still forming turbid and transient plaques.

**Figure 5.**
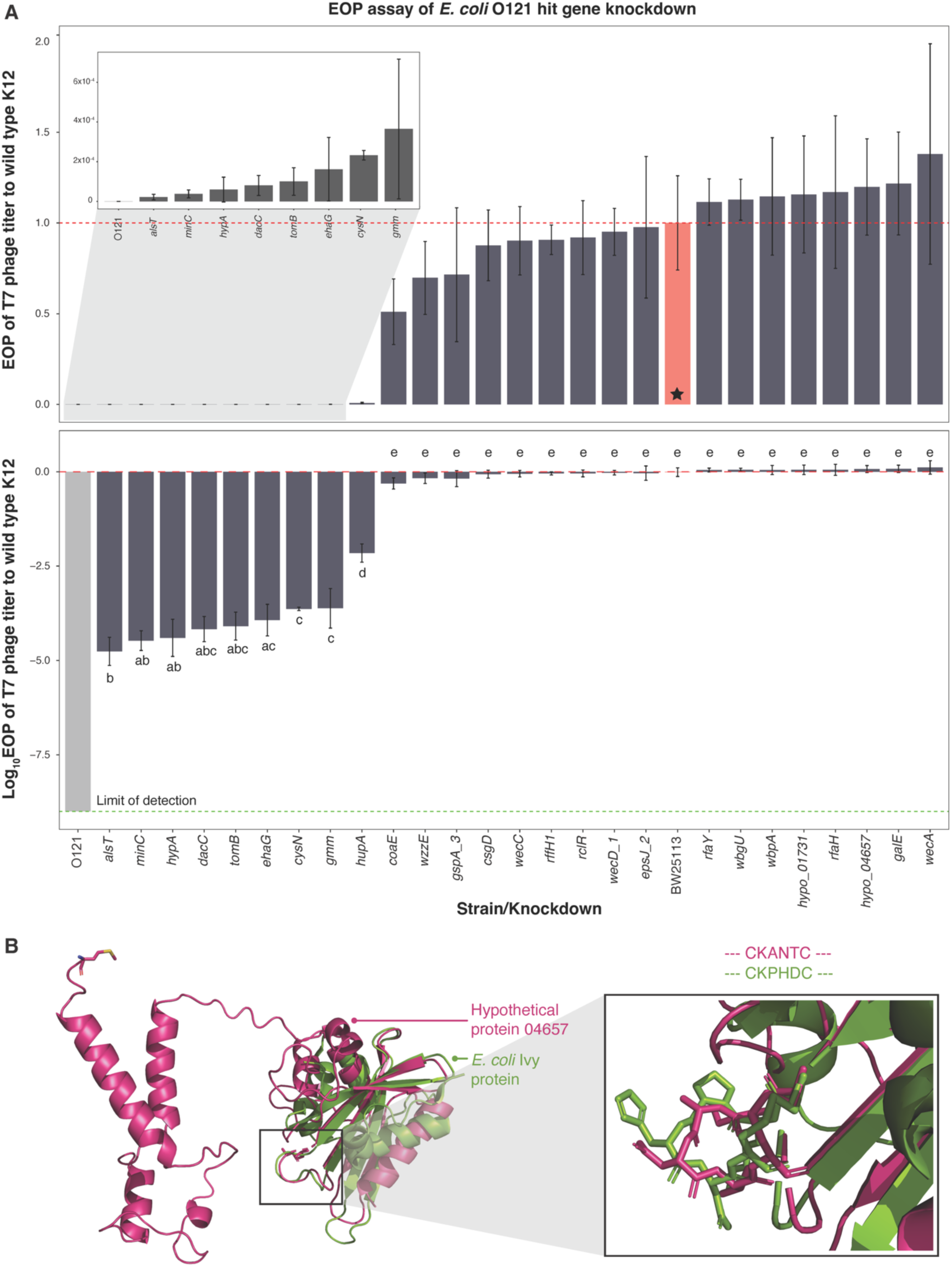
Efficiency of plating confirms the diverse impact of the identified factors on phage. (A) We selected 26 genes identified in our screen of T7 phage on *E. coli* O121 to validate the effect of individual gene knockdown by efficiency of plating (EOP). EOP was quantified in each clonal gene knockdown on *E. coli* O121 with comparisons made to either *E. coli* BW25113 or O121 containing *rfp* sgRNA as a control. EOP values are shown top without transformation to facilitate comparison, where an EOP of 1 is equivalent to plaque capability on permissive BW25113 (pink bar and a horizontal dashed line, starred). EOP values are shown bottom as log_10_ transformed. All data are reported as the mean EOP±SD across biological replicates. Difference in EOP was determined by Tukey’s HSD test at the *p-*value of 0.05. (B) ColabFold (AlphaFold2) was employed to predict the protein structure of Hypo_04657 (depicted in pink). This protein comprises two significant domains: the C-terminal domain characterized by *α*-helices (on the left) and the N-terminal domain featuring both *α*-helices and β-sheets (on the right). The right domain was superimposed onto the monomeric structure of the *E. coli* Ivy protein (PDB: 1XS0, shown in green), demonstrating a root mean square deviation (RMSD) of 0.81 Å. A close-up view highlights the conserved inhibitory loop “CKPHDC” in the *E. coli* Ivy protein, contrasting with the “CKANTC” loop observed in the Hypo_04657 protein.

In *E. coli* O121, approximately a thousand hypothetical proteins lack a well-established function. We found that knocking down three of these hypothetical proteins, 01731, 04657, and 05030 (*hypo_01731*, *hypo_04657*, *hypo_05030*), increased phage fitness. We predicted the structure of these proteins using ColabFold(55, 56) and predicted protein function using DeepFRI(57). The *hypo_01731* gene is located between *rho* and *wec* operons (Supplementary Figure S11A). Thus, given the significant improvement of efficiency of plating, we hypothesized its involvement in the outer structure, specifically in ECA and O-antigen. Similarly, knockdown of *hypo_04657* significantly improves plaquing efficiency; therefore, this gene is potentially relevant to the bacterial surface.

We aimed to gain more insight about *hypo_04657*, which is surrounded by other hypothetical genes. The N-terminal half of *hypo_04657* is predicted to have hydrolase activity due to structural similarity to the inhibitor of vertebrate lysozyme (Ivy; PBD:1XS0) in *E. coli*. Structural alignment of the Ivy*-*like region of Hypo_04657 and *E. coli* Ivy monomer showed a distance of 0.81Å. Additionally, *hypo_04657* has“**CK**ANT**C**” instead of the conserved “CKPHDC” motif(58) (Figure 5B). The C-terminus of Hypo_04657 is a putative *α*-helical transmembrane domain. We reasoned that Hypo_04657 may act as an inhibitor of lysozyme, potentially with higher specificity towards phage lysozyme. Given that the T7 phage produces bifunctional lysozyme, which serves to both lyse the cell wall and inhibit T7 transcription, knocking down *hypo_04657* could improve phage replication and the release of phage progenies(59, 60). Lastly, the Hypo_05030 was structurally similar to the RelE/ParE type II TA system found in γ-Proteobacteria (Supplementary Figure S11B and S11C). These toxins inhibit translation, leading to a stalling or cleavage of mRNA. We anticipated that Hypo_05030 may reduce the translation of both phage and bacterial mRNA during the infection, ultimately resulting in a decrease in phage reproduction(61, 62).

### The impact of host genes on phage fitness shapes phage evolution

Few studies have investigated phage-host co-evolution from a perspective other than host defense and phage anti-defense systems. Given that phages co-evolve with their hosts to outcompete each other and other community members(63), we hypothesized that our experimental data of host-influenced phage fitness should reflect general principles of phage evolution, representing the impact of host factors beyond just the replication step.

We hypothesized that the host genes conferring a beneficial impact on phages would likely be acquired by phages over the course of evolution, while host genes detrimental to phages would not be acquired. To explore this hypothesis, we cataloged the prevalence of *E. coli* K-12 and B strain genes in a curated subset of the IMG/VR v4(64) viral genome database consisting of only high-quality, dereplicated viruses (Supplementary Figure S13A). Additionally, we ignored *E. coli* genes that were likely to belong to integrated prophages (Supplementary Figure S13B). As expected, host genes that supported phage fitness in both permissive strains were highly abundant in viral genomes (Figure 6A, Table S7) and statistically enriched compared to *E. coli* genes that did not affect phage fitness (Figure 6B, Mann-Whitney U test, *p*-value<0.05). The simplest explanation is that since these genes are necessary for optimal phage reproduction, many other viruses have acquired these genes to account for the potential absence of these genes among different host strains.

**Figure 6.**
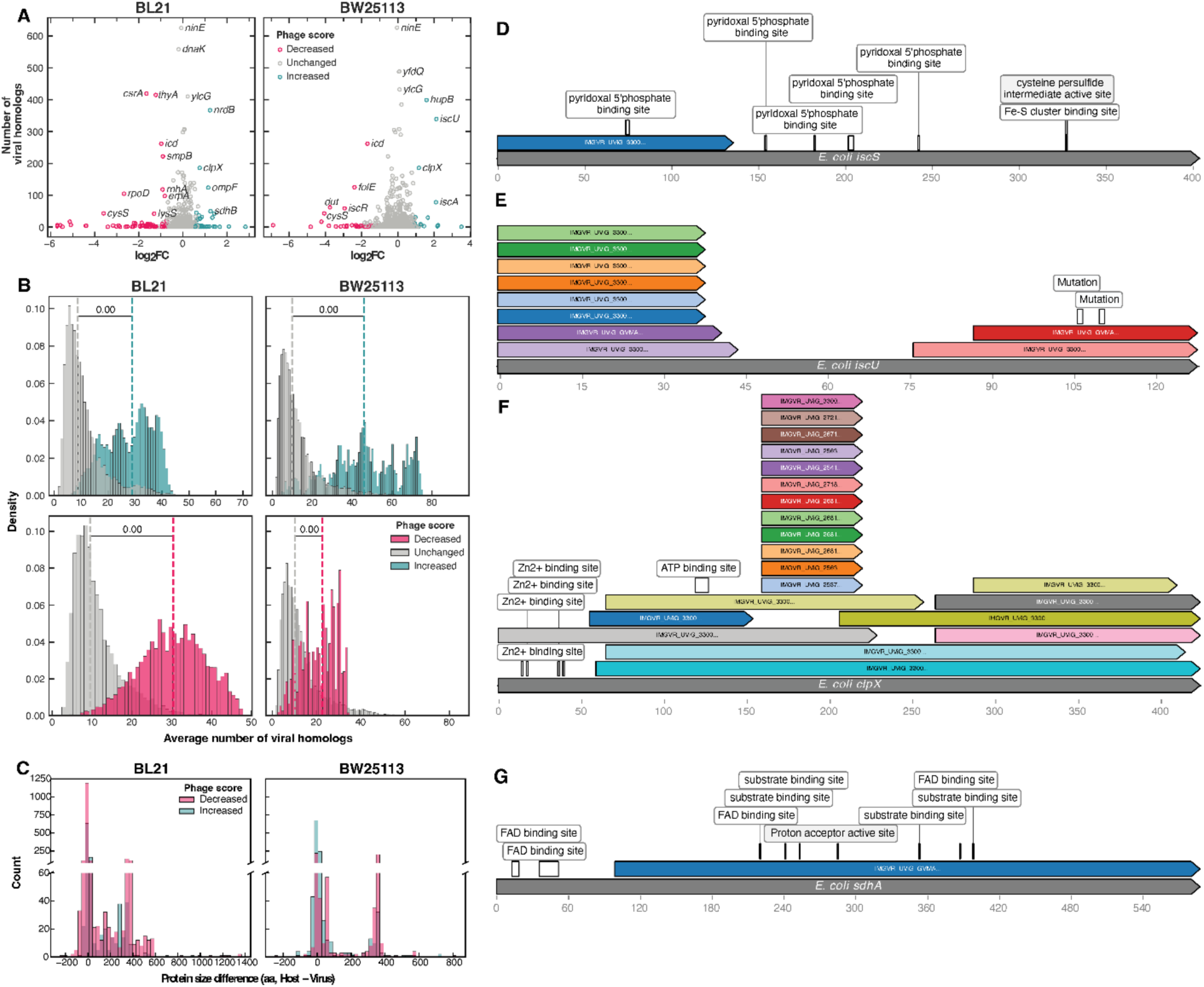
Viruses carry host genes that both promote and inhibit viral fitness. (A) Experimentally determined log_2_ fold change values upon CRISPRi knockdown compared to the abundance of each gene in a curated subset of the IMG/VR v4 database. The subset included 249,027 high-quality viral operational taxonomic unit representatives, corresponding to 13,363,675 proteins. Hits were determined using a minimum criterion of an e-value≤10^-5^ and sequence identity≥50% over≥50% of the viral protein. (B) Gene sets of size equal to 60% of the number of genes where experimental treatment led to an increase or decrease in phage fitness were randomly sampled 10,000 times. The distribution of the average number of viral homologs from each gene set is shown, comparing samples only consisting of the significant genes to samples only containing genes that did not affect phage fitness. Dotted vertical lines represent the median of each distribution, and the value corresponds to the *p*-value of a Mann-Whitney U test. (C) For each virus-host homolog pair, the distribution of the difference in protein length in amino acids is shown. Values above 0 indicate the host homolog is longer, while values below 0 indicate the viral homolog is longer. (A–C) Compare results from *E. coli* BL21 and BW25113 strains. Data are colored based on whether CRISPRi knockdown of gene did not affect (grey, “unchanged”), increased (green), or decreased (red) phage fitness. (D– G) Four case studies of genes carried in viruses whose CRISPRi knockdown led to an increase phage fitness in at least one *E. coli* strain tested: (D) *iscS*, (E) *iscU*, (F) *clpX*, and (G) *sdhA*. The bottom bar represents the full-length *E. coli* protein, and the colorful upper bars are full-length viral proteins. The white or gray annotations mark important sites in the *E. coli* protein per the Uniprot database.

Interestingly, although we predicted that host genes detrimental to phage fitness would not be abundant across viral genomes, these host genes were, in fact, also highly abundant in viral genomes (Figure 6A) and statistically enriched (Figure 6B, Mann-Whitney U test, *p*-value<0.05). While the acquisition of these genes could confer benefits to phages under other conditions or on other bacterial hosts, we propose a more likely hypothesis, the dominant-negative effect (DNE). This phenomenon involves genetic mutations or truncations that adversely alter the function of wildtype proteins by binding to or competing with their wildtype homolog. To explore our hypothesis, we examined the difference in protein sequence length for each host-phage homolog pair. While most homolog pairs showed no difference in length, many host homologs were significantly larger than the corresponding viral homolog (Figure 6C). This indicates that viruses carry truncated copies of host genes. These truncated variants could inactivate the corresponding host proteins by poisoning the pool with nonfunctional proteins as a DNE. Additionally, evidence of DNE has been identified before in lambda-like phage lysis regulation(65). Notably, cases include truncated versions of *clpX*, *iscS*, *iscU*, and *sdhA* that likely partially or fully inactivate the protein (Figure 6D-6G). For example, certain viral homologs of the *clpX* are missing the Zn-binding domain, which is crucial for ClpXP activity(66). A viral *iscS* copy is missing the active site and coenzyme binding sites, while *iscU* variants have mutations in sites that lead to slower Fe-S cluster assembly(67). Further, these viral *sdhA* genes lack the N-terminus that composes most of the FAD binding site, likely leading to reduced succinate dehydrogenase activity. These findings supported that DNE is one of the widespread mechanisms for phage to manipulate the host intracellular state and optimize infection conditions.

## Conclusions

Our comprehensive investigation using PHAGEPACK has unveiled new insights into bacterial factors influencing T7 phage infection. Unlike traditional host-based screens(5, 6, 21), PHAGEPACK directly quantifies phage fitness rather than host fitness. We identified previously unknown *E. coli* pathways affecting T7 infectivity, including iron-sulfur cluster biogenesis, polyadenylation and post-transcriptional processes, codon usage, and TA systems. Furthermore, our study of the non-permissive pathogen *E. coli* O121 highlighted Enterobacter common antigen biosynthesis, O-antigen biosynthesis, and inner core modifications of the bacterial membrane as major impediments to phage infection and replication. PHAGEPACK also revealed dozens of genes with moderate effect sizes, including those involved in transcriptional regulation, RNA synthesis, and glycerol metabolism. The strength of PHAGEPACK lies in its systematic quantification of each host factor’s impact on phage fitness.

There are potential sources of false positives and false negatives in PHAGEPACK. One significant limitation that could lead to false negatives is the variability in CRISPRi repression efficiency across the *E. coli* genome, which has been shown to vary up to 10-fold in previous studies(68). Genes that show leaky expression despite CRISPRi repression might result in false negatives. To mitigate this, we use four sgRNAs per gene. Another assumption we make is that phage packaging correlates with phage fitness. However, there may be cases where a gene knockdown affects phage packaging without impacting phage fitness, leading to false positives. Our limited clonal validation of 26 genes in *E. coli* O121 showed no false positives. Without knowledge of which host genes specifically influence phage packaging, we cannot preemptively eliminate such false positives, should they exist.

We discovered that homologs of host factors beneficial and detrimental to phage fitness can be found on viral genomes. Viruses inherently possess limited genome capacity; therefore, any genes obtained and retained likely provide benefits under certain conditions. It is intriguing that phages carry homologs of genes that are detrimental to phage fitness. One plausible reason could be the dominant negative effect (DNE). Generally, DNE occurs when a mutated protein interferes with the function of wildtype protein, causing a loss-of-function in protein. In eukaryotes, this phenomenon has implications for various genetic disorders and diseases. For instance, the mutant of the p53 tumor suppressor protein can lead to an inhibition of wildtype p53, contributing to the development of oncogenesis within the cell(69). Further, a study utilizing high throughput sequencing of yeast peptide libraries has elucidated the impact of DNE on cellular growth. This study has revealed that DNE can affect a diverse array of cellular components involved in processes such as cell cycle, stress response, and translation, highlighting the extensive role of DNE in modulating cellular processes(70). In a similar vein, carrying host factor homologs may be a viral strategy to counteract host defenses and tailor host metabolic and physiological states for optimal infection conditions. This phenomenon remains unexplored in the context of bacteria-phage interactions and is an exciting prospect for therapeutic applications.

PHAGEPACK can be generalized because packaging sequences can be identified in many phages and CRISPRi has been implemented in many bacterial hosts. Phage packaging sequences can be identified using either the standard ligation-digestion method(71) or NGS-related PhageTerm predictor(72), a tool designed to detect DNA termini and predict phage packaging sequences in dsDNA phages based on read pattern and bias observed in NGS data of the phage genome. This tool utilizes the common location of phage packaging recognition sequences at the DNA termini, although the sequences may differ across different phages. Packaging sequences and schemes have been characterized in various phages; for instance, λ and HK97 phages rely on *cos* sequence, T3 and T7 phages use DTR, and P1, P22, SPP1, and T4 rely on the *pac* site. PhageTerm has been shown to successfully determine the phage termini and packaging scheme of uncharacterized *Clostridium difficile* prophages, such as phiCD211, phiCD481, phiCD506, and phiMMP01. With the versatility of CRISPRi knockdown in diverse bacterial hosts(73, 74) and the availability of these tools, PHAGEPACK can broaden our investigation across various phage-bacteria pairs.

Phage therapy efforts frequently encounter challenges due to the development of host resistance to phages. Typically, when resistance arises, phage biologists search for alternative phages. We see PHAGEPACK as a valuable tool for uncovering the genetic mechanisms by which bacteria develop phage resistance, offering critical insights for developing strategies to combat bacterial resistance. By applying PHAGEPACK to various strain variations of a pathogen, we can identify vulnerabilities common to the species, which could be leveraged to create broadly effective phages. Additionally, using PHAGEPACK under different environmental conditions—such as varying nutrients, salt levels, and pH—will uncover the conditional effects of host genes, guiding the fine-tuning of phage activity through environmental adjustments. In summary, PHAGEPACK provides a framework for gaining insights into bacterial resistance, developing precisely tailored phage therapy strategies, and maximizing the therapeutic potential of well-characterized phages that are currently available, thereby advancing their clinical utilization.

## Materials and Methods

### Bacterial strains, bacteriophage, and culture conditions

The T7 bacteriophage was obtained from ATCC (ATCC BAA-1025-B2). *E. coli* 10G, a highly competent DH10β derivatives originally obtained from Lucigen (60107–1)(75), and *E. coli* BL21, a laboratory stock, were used. *E. coli* BW25113, obtained from Douglas Weibel (University of Wisconsin – Madison), and its derivatives, including BW25113Δ*trxA* and other BW25113 single gene knockouts, were derived from Keio collection(76). *E. coli* O121 was obtained from ATCC (ATCC BAA-2190), and *E. coli* BL21 (REL606) was obtained from Robert Landick (University of Wisconsin – Madison). Additionally, *E. coli* BW25141 *pir*^+^ carrying pJMP1039, pJMP1189, and pJMP1339 were obtained from Jason Peters (University of Wisconsin – Madison).

All bacterial cultures were grown in LB media (1% tryptone, 0.5% yeast extract, and 1% NaCl) and plated on LB agar plate (LB media with addition of 1.5% agar). LB media was consistently used for all experiments. Carbenicillin (100 µg/ml final concentration for all sgRNA delivering plasmids and pJMP1039), Kanamycin (50 µg/ml final concentration for pJMP1189 and Keio single gene knockout strains), or combinations as required were added for selection. Isopropyl β-D-1-thiogalactopyranoside (IPTG, 1mM final concentration) was added to induce CRISPRi. Incubation of all bacterial cultures occurred at 37°C with liquid cultures subjected to shaking at 200-250 rpm. Bacterial hosts were streaked on appropriate LB plates, incubated, and stored at 4°C.

The T7 bacteriophage was originally propagated using *E. coli* BL21 upon receipt from ATCC. Subsequently, propagation was carried out using *E. coli* 10G and BW25113 for all phage selection experiments. These experiments were conducted using LB media and followed the bacterial culture conditions described for bacterial hosts. Phages were stored in LB at 4°C.

All bacterial hosts were stored as liquid samples at −80°C in 25% glycerol and 75% relevant media for long-term storage.

SOC (2% tryptone, 0.5% yeast extract, 0.2% 5M NaCl, 0.25% 1M KCl, 1% 1M MgCl_2_, 1% 1M MgSO_4_, 2% 1M glucose in dH_2_O) was used for the recovery of bacterial hosts after transformations. Diaminopimelic acid (DAP) was supplemented at 0.3mM to support growth of *dap-*negative *E. coli* strains.

#### General cloning procedure

PCR was performed using KAPA HiFi (Roche KK2101) for all experiments, except for colony PCR, where KAPA2G Robust (Roche KK5005) was utilized. The New England Biosciences (NEB) Golden Gate Assembly Kit (BsaI-HF v2, E1601L) and the Gibson Assembly Protocol (NEB E5510) were employed, utilizing our in-house-prepared Gibson Assembly Master Mix (final concentration 100mM Tris-HCl pH 7.5, 20mM MgCl_2_, 0.2mM dATP/dCTP/dGTP/dTTP, 10mM DTT, 5% PEG-8000, 1mM NAD^+^, 4U/ml T5 exonuclease, 4U/µl Taq DNA ligase, 25U/ml Phusion polymerase). Restriction enzymes were obtained from NEB, except for DNase I (Roche 4716728001). DNA purification utilized E.Z.N.A® Cycle Pure Kits (Omega Bio-tek D6492-01) with a centrifugation protocol, and DNA was concentrated using DNA concentrator (Zymo Research D4004). Gel extraction was performed using E.Z.N.A® Gel Extraction Kit (Omega Bio-tek D2500-01) with a centrifugation protocol. All cloning procedures followed the manufacturer’s documentation, with specific conditions and/or variables noted if different from the standard protocol.

PCR reactions used 0.1 ng of plasmid or 1 µl of phage stock (with an approximate titer of 10^9^ PFU/ml) directly as a template. Standard reaction cycle was applied, involving initial denaturation at 95°C for 3 minutes, denaturation at 98°C for 20 seconds, extension and final extension at 72°C, and annealing at temperature and duration relevant to primers and targeted amplification size. Phage genome fragments were amplified directly from the phage as a template without genome extraction, employing a modified PCR protocol with an initial denaturation at 95°C for 5 minutes. PCR product was purified using a DNA purification kit (Omega Bio-tek) and a Spin Column for DNA (EconoSpin 1920-250), following the manufacturer’s documentation with CP Buffer (Omega Bio-tek PDR042) and DNA Wash Buffer (Omega Bio-tek PDR044).

DpnI digestion was conducted on all PCR reactions using plasmid templates. The digestion was performed directly on the PCR product immediately before purification. For each 20 µl PCR reaction, 10 units of DpnI were mixed with 5 µl of 10x CutSmart Buffer, and dH_2_O was added to reach a total volume of 50 µl. The reaction was incubated at 37°C for 2 hours and subsequently heat-inactivated at 80°C for 20 minutes.

DNase treatment was applied on the phage solution obtained from the phage selection experiment. This involved combining 10 µl of precipitated phages, 2 µl of 10U/µl DNase I, 4 µl 10x DNase buffer, and dH_2_O to a total volume of 40 µl. The mixture was then incubated at 37°C for 1 hour, and subsequently inactivated at 75°C for 15 minutes. The reaction was purified to obtain plasmid originally packaged in the phage during the selection experiment, using column purification kit, and eluted with TE Buffer (1mM EDTA, 10mM Tris-Cl, pH 8.0) or water. Depending on the number of plasmids packaged, a DNA concentrator may be used to increase the final concentration of purified plasmid DNA during PCR purification.

Transformation was carried out through electroporation using a Bio-rad MicroPulser (165–2100). The setting used were Ec1 (1-mm cuvette, 1.8 kV) for plasmid transformation and Ec2 (2-mm cuvette, 2.5 kV) for CRISPRi sgRNA library transformation. For transformation, we employed 30-100 µl of competent cells and 1-2 µl of DNA, except for 10-20 µl of dialyzed Golden Gate Assembly for sgRNA library transformation in cloning strain *E. coli* 10G. To ensure the sufficient coverage, we utilized 100-400 ng of sgRNA plasmid library for each transformation in the target bacterial host. After electroporation, bacterial cells were immediately recovered by adding 950 µl pre-warmed SOC, followed by a recovery at 37°C for 45 minutes to 1 hour, and then plated on relevant media.

*E. coli* competent cells, including 10G, BL21, BW25113, and O121, were prepared by inoculating 4 ml of overnight culture into 196 ml SOC (1:50 dilution) with antibiotics as needed. Our competent cells were initially grown at 21°C at 200 rpm, reaching an OD_600_ of approximately 0.4-0.6, determined using an Agilent Cary 60 UV-Vis Spectrometer and Ultrospec 10 Cell Density Meter (Amersham Biosciences). Some competent cells were alternatively grown at 30°C with comparable transformation efficiency. Following centrifugation at 4°C (800-1500xg for 20 minutes), the supernatant was discarded, and 40 ml of 10% cold glycerol was added. Bacterial cells were gently homogenized and washed. After three washes, the final cells were resuspended in approximately 1 ml of 10% cold glycerol, aliquoted, and stored at −80°C until needed for transformation. These cells were electrocompetent for plasmid library transformation.

DNA quantification was performed using NanoDrop 2000 (Thermo Scientific) with 1 µl of DNA, except for Next Generation Sequencing DNA, which was quantified using Qubit 4 fluorometer (Invitrogen) following the manufacturer’s documentation.

Detailed protocols for cloning, as well as a list of all primers used in the experiments presented in this publication, are available on request and provided in Table S1.

#### Plasmid description

pJMP1039, pJMP1189, and pJMP1339 were provided by Jason Peters (University of Wisconsin-Madison). All three plasmids, featuring R6K gamma origin of replication, require *pir* gene for replication (BW25141 *pir*^+^ strain). pJMP1039 served as a helper plasmid, carrying *transposaseABCD* gene and pJMP1189 functioned as a donor plasmid, containing *Hsa Spy dCas9* gene and *mrfp* gene. The *dCas9* gene was expressed under inducible *lac* promoter.

pUC19, purchased from NEB, was served as the backbone for the all sgRNA delivery plasmids. The original BsaI site presented on ampicillin-resistant (*ampR*) gene was eliminated through site-directed mutagenesis.

pT7cand01, pT7cand02, pT7cand03, and pT7cand04 were constructed as pUC19 vectors containing T7 packaging sequences (candidate 1-4, as illustrated in Figure 1B) and *ampR* gene.

psgRNA01 is a pUC19 vector containing sgRNA inserted, *lacI* gene, and *ampR.* This plasmid was generated by removing the T7 packaging sequence (candidate 3) out from pT7sgRNA01.

pT7sgRNA01 is a pUC19 vector containing T7 packaging sequences (candidate 3), *lacI* and *lacIq* promoter, *ampR* gene, sgRNA region and gRNA scaffold. The sgRNA is expressed under an inducible *lac* promoter. This plasmid is designed for the delivery of sgRNA in CRISPRi pooled experiments.

All plasmid and individual sgRNA sequences used in the individual gene knockdown experiments are listed in Table S1.

#### Phage packaging quantification

Phage genome packaging fragments were amplified by PCR as described earlier. Each packaging candidate was cloned into pUC19 backbone. The plasmid was first transformed into the cloning *E. coli* 10G and then target bacteria strains, and the transformants were selected on selective media. The bacterial cells carrying the phage packaging plasmid were grown overnight, back diluted (1/50 dilution), and then cultured to reach the exponential growth phase estimated by OD_600_∼0.6. Wildtype T7 phage was mixed with the bacterial culture at an MOI of approximately 2 and incubated for 1 hour for an infection. The culture was filtered using a PES 0.22 µm filter (Cell Treat 229747) to collect phage lysate.

Phage packaging quantification was assessed through a transduction experiment using *E. coli* BW25113Δ*trxA* as the recipient, a strain that T7 phage cannot replicate properly due to a lack of thioredoxin A facilitating T7 DNA polymerase during plaque formation. This strain was obtained from the Keio collection. *E. coli* BW25113Δ*trxA*::*kanR* was cultured overnight with kanamycin-selective media (final concentration 50 µg/ml) and back-diluted to exponential phase on the day of the experiment. Phage lysate was subjected to a 10-fold dilution (from 10^0^ to 10^-7^ dilution), and 15 µl of each diluted phage was mixed with 150 µl of *E. coli* BW25113Δ*trxA*::*kanR.* The mixture was incubated at 37°C with shaking at 200-250 rpm for 40 minutes. Subsequently, 10 µl for each phage dilution mixture was dropped in triplicate onto carbenicillin and kanamycin (Crb-Km) selective agar media (final concentration 100 µg/ml carbenicillin and 50 µg/ml kanamycin) and incubated overnight at 37°C. The number of phages containing packaged plasmid is indirectly quantified through the number of recipients that is transduced and grown on selective media containing both Crb-Km. Number of transduced recipients was determined by counting the colony-forming unit (CFU/ml) of recipient cells, providing the absolute number of plasmids packaged phages produced. Statistical analysis was performed by Tukey’s honestly significant difference (HSD)(77) at the *p-*value of 0.05.

#### *dCas9* integration by co-transformation and conjugation

The bacterial host was engineered with the integration of the *dCas9* gene into the genome using a standard transformation and mobile CRISPRi method(73). Initially, we co-transformed pJMP1039 (*transposaseABCD* enzyme) and pJMP1189 (*dCas9* gene, *mrfp*, *kanR*) into electrocompetent *E. coli* BW25113 and BL21 cells, selected transformants on kanamycin-selective agar media. Due to low efficiency with co-transformation, a conjugation suggested in the original protocol(73) was performed to integrate the *dCas9* gene into the clinically relevant *E. coli* O121 strain.

In brief, cultures of the donor *E. coli* BW25141 (harboring pJMP1039 and pJMP1189) were grown overnight in LB with DAP (0.3mM), and the recipient *E. coli* O121 was cultured in LB. After gently spinning down 1 ml of cells at 3000 rpm for 5 minutes, we washed them with PBS (137mM NaCl, 2.7mM KCl, 10mM Na_2_HPO_4_, 1.8mM KH_2_PO_4_), and resuspended the cells in 500 µl PBS. We measured OD_600_ to estimate the cell concentration and combined the donor and recipient cells in a 1:1 ratio, mixed by pipetting twice, plated 100 µl of the mixture onto non-selective agar plate containing DAP without spreading, and incubated the plate overnight at 37°C, 24 hours (conjugation times can vary). The conjugation mixture was scraped up, mixed with 1 ml PBS, vortexed (Southwest Science SBV1000), gently spun down, and the wash step was repeated. The conjugation mixture was resuspended in 1 ml PBS, and 100 µl of both undiluted and 10-fold diluted mixtures were plated onto selective plate (Km, DAP^-^). Single colonies were picked, re-streaked on a selective agar plate, and verified via colony PCR of *dCas9* gene insertion.

#### CRISPRi library sgRNA spacer design and construction

We designed sgRNA spacers using the protocol described by Amy *et. al.*(74), and the sgRNA_design script can be downloaded from https://github.com/ryandward/sgrna_design. Bacterial genomes, including *E. coli* K12 (NC000913.3), BL21 (CP000819.1), T7 phage genome (V01146.1), and *E. coli* O121 BAA-2190 (genome sequences available on the ATCC website), were utilized to design targeted sgRNA spacers. The script designed 20-bp length sgRNA spacers by identifying PAM site (NGG) regions on bacterial genomes and provided sgRNA spacer sequence, target region, and specificity. We selected four non-overlapping sgRNA spacers, unless the gene was small and required overlapping sgRNAs or fewer than four sgRNA spacers. All sgRNAs were selected with the highest specificity score (SPECIFICITY==39) and designed to target the non-template strand (as antisense or “anti”) to maximize knockdown efficacy (see Table S2). As a control, we included 1,000 randomly generated sgRNA spacers. Two sets of sgRNA spacer libraries were designed for *E. coli* K12/B (total of 23,581 sgRNAs) and *E. coli* O121 (total of 19,583). For this study, we employed 19,471 sgRNAs specifically designed for use in *E. coli* K12 and B strains, excluding the 4,110 sgRNAs intended for an unrelated study.

We purchased a pilot sgRNA library from Integrated DNA Technologies (IDT), which consisted of a total of 51 sgRNA spacers (36 sgRNAs targeting 9 specific genes and 15 randomized sgRNAs, Table S2). The library was resuspended to 100µM single-stranded DNA (ssDNA). To generate double-stranded DNA (dsDNA) oligos, we performed PCR using a single primer (initial denaturation at 95°C for 3 minutes, annealing at 60°C with a ramp of 0.1s/°C for 15s, extension at 72°C for 20s). The PCR product was diluted by one-third, and 1 µl was used as a template for sgRNA oligo amplification (total of 25 cycles). Subsequently, we assembled the pilot sgRNA library using the Golden Gate Assembly Kit, dialyzed the product using 0.025 µm MCE Membrane (MF-Millipore VSWP02500), transformed it into *E. coli* 10G, quantified the number of transformants on Crb and Km selective agar plates, and grew transformants in Crb-Km selective liquid media at 37°C overnight.

The sgRNA library for *E. coli* K12 and B strains was obtained from Jason Peters (University of Wisconsin – Madison) and then amplified to dsDNA sgRNA oligos. A pool of 0.01 pmol sgRNA oligos was used as a template for PCR (with initial denaturation at 95°C for 3 minutes, annealing at 60°C with a ramp rate of 0.1s/°C for 15 s, and extension at 72°C for 20 s). Oligos were further extended through PCR to accommodate size limitations encountered in column purification. The sgRNA library was assembled using Golden Gate assembly at a 1:2 molar ratio of pT7sgRNA01 backbone to sgRNA spacers, with long cycling conditions ((37°C for 5 minutes, then 16°C for 5 minutes) x 30 cycles, followed by 60°C for 5 minutes). The resulting library was dialyzed using 0.025 µm MCE Membrane (MF-Millipore VSWP02500), transformed into *E. coli* 10G, and cultured in Crb-Km liquid media (∼1ml of transformants in 50-200 ml media). Transformants were selected and counted on selective agar plate immediately after a transformation to quantify the number of transformants and assess library coverage. The sgRNA library was extracted, transformed (100-200 ng) to *E. coli* BW25113 and BL21, selected and counted Crb-Km agar plate, and cultured in Crb-Km liquid media at 37°C. The concentration of sgRNA library in CFU/ml was quantified from agar plates after overnight incubation.

The sgRNA oligo pool targeting *E. coli* O121 genome was purchased from Twist Bioscience (San Francisco, CA) and resuspended in 10mM Tris buffer to a concentration of 10 ng/µl as recommended in the documentation. We amplified sgRNA oligos using specific primers in 12 cycles of PCR using 5 ng of template, purified the product, and assembled it with pT7sgRNA01 backbone using Golden Gate Assembly Kit. After dialysis, the assembly was transformed into *E. coli* 10G, transformants were quantified on Crb-Km agar plates, and the culture was grown overnight at 37°C in selective liquid media.

All pT7sgRNA01 libraries were first transformed into *E. coli* 10G. We extracted the library plasmid from overnight transformants and transformed into the target *E. coli* host strains. Transformants were quantified on Crb-Km selective agar plates, grown overnight in selective liquid media, collected, and stored as glycerol stocks in −80°C.

The library coverage was validated by deep sequencing at both the initial transformation in *E. coli* 10G (Supplementary Figure S2A and S6) and in the targeted strains.

#### RFP production quantification by plate reader experiment

We selected sgRNA spacer targeting *mrfp* gene with the highest repression activity(73) (see Table S1), cloned it into pT7sgRNA01 using Golden Gate Assembly, and transformed the plasmid into *E. coli* BW25113, BL21, and O121. The transformants were selected on Crb-Km agar plate, and the construction was verified using colony PCR.

We inoculated bacterial culture both with and without the induction of CRISPRi, achieved by adding 1mM IPTG directly to liquid media (Crb-Km), and incubated them overnight at 37°C. For the plate reader experiment, the overnight bacterial culture was diluted 1:10,000 in fresh LB media with and without the inducer (1mM IPTG). Next, 200 µl aliquots were dispensed into 96-well tissue culture plate (Cell Treat 229195) in triplicates, including the original bacterial host as a control (wildtype and lacking pT7sgRNA01). The experiment was conducted using a BioTek Synergy HTX Multi-Mode Microplate reader with Gen 5 software (Agilent), set at a temperature of 37°C, continuous linear shaking, and OD measurement at 600 nm. Relative Fluorescence Units (RFU) were measured with excitation at 540/35 and emission at 600/40, and data were collected every 15 minutes. RFU values were normalized by OD_600_, and the average relative RFU was calculated. The relative amount of PFR produced under induced and non-induced states was determined where the relative RFUs were consistent (Supplementary Figure S1A). The difference of relative RFU production between induced and non-induced condition was determined using Tukey’s honestly significant difference (HSD)(77) at the *p-*value of 0.05.

#### Gene expression quantification by RT-qPCR

We constructed pT7sgRNA01 vectors targeting three chosen genes (*lpcA, trxA, waaC*) using Golden Gate Assembly Kit and cloned them into *E. coli* K12. The bacterial clones were cultured overnight with CRISPRi induction (1mM IPTG). After back-dilution (1:50) and growth to an exponential phase (OD_600_∼0.5-0.6), we performed RT-qPCR by collecting 1 ml of bacterial culture. The cells were centrifuged at 10,000xg for 1 minute, supernatant was discarded, and the culture was stored at −80°C, when immediate RNA extraction was not possible. We extracted RNA using TRIzol™ Reagent (Invitrogen™ 15596026). Briefly, 1 ml TRIzol was added to the bacterial culture, vortexed for 1 minute, and incubated at room temperature (RT) for 5 minutes. Following, 200 µl pre-chilled chloroform (0.2x volume of TRIzol) was added, gently inverted for 15 seconds, incubated at RT for 3 minutes, and centrifuged at 12,000xg for 15 minutes at 4°C. The aqueous phase, without disturbing the interphase, was transferred to a new tube and subjected to cleanup using RNA cleanup and concentrator kit (Zymo R1016). This involved adding 2x volume of RNA binding buffer and an equal volume of ethanol, and gently pipetting for mixing. Approximately 700-800 µl of the mixture was transferred to a column-IC (Zymo C1004-5), washed twice with 700 µl and 400 µl RNA wash buffer for the first and second wash, respectively, and centrifuged at 15,000xg for 30 seconds. The flow-through was discarded after each wash, and the column was spun dry for 2 minutes before the final elution with 15 µl RNase-free water.

DNase I digestion was performed using DNase I (Roche 4716728001). We mixed 15 µl of the RNA sample with 20U of DNase I in 50-µl reaction, incubated at 37°C for 30 minutes, stored at 4°C without heat inactivation. The digested sample was purified using RNA cleanup and concentrator kit (Zymo R1016), eluted with 15 µl RNase-free water. RNA was quantified using NanoDrop 2000 (Thermo Scientific) and proper purity was verified with A260/A280∼2.00.

RT-qPCR was conducted using Luna®Universal One-Step RT-qPCR Kit (NEB E3005L) following the manufacturer’s documentation. Primer efficiency was validated using RNA samples from *E. coli* without gene knockdown, ranging from 1 to 0.001 ng/µl. For each reaction, 1µl of RNA template was added to a 9 µl Master Mix (5 µl Luna Universal One-Step Reaction Mix, with and without 0.5 µl Luna WarmStart RT Enzyme Mix, 0.4 µl 10µM Forward primer, 0.4 µl 10µM Reverse primer, dH_2_O up to 9 µl). The reaction was performed in MicroAmp® Fast Optical 96-Well Reaction Plate with Barcode 0.1 ml (Applied Biosciences 4346906) and run in 7500 Real-Time PCR system (ThermoFisher Scientific). Primers with efficiency (>90%, Supplementary Figure S14A-S14D) were employed for each target gene RNA quantification. For each target gene quantification, 20 ng total RNA extracted from bacterial culture (*lpcA, trxA, waaC*) with and without IPTG-induced gene knockdown was used as a template. The housekeeping gene target was *rrnA* (Supplementary Figure S1B).

Primer efficiency was calculated using the slope obtained from the linear equation of C_t_ and RNA concentration.

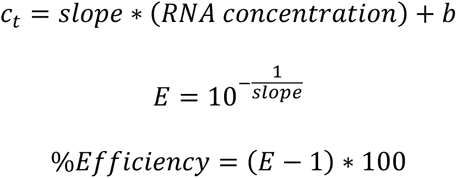

Gene expression was determined using ΔΔC_t_ calculated using these following equations.

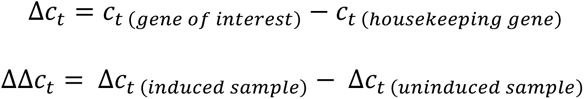

#### Phage selection and CRISPRi experiment

Bacterial cultures carrying the sgRNA library and CRISPRi system were inoculated from glycerol stock into selective liquid media (Crb-Km and 1mM IPTG), incubated at 37°C overnight, and back-diluted (1:50) on the day of the experiment to OD_600_∼0.5. Two milliliters of the bacterial culture were aliquoted into two culture tubes for phage selection and control conditions. Phage, at relevant MOI (phage diluted if needed), was added with the estimation that OD_600_ of 1 is approximately equal to 8×10^8^ CFU/ml (Agilent biocalculator). The MOI of 0.01 was administered to *E. coli* K12 and B strains, whereas the MOI of 2 was applied to *E. coli* O121 ensuring sufficient phage encounters with at least one host cell on this non-permissive host. The mixture was incubated at 37°C for 30 minutes (*E. coli* K12 and B strains), 45 minutes and 2 hours (*E. coli* O121). The latter was performed at two timepoints, taking into account uncertainties in the latency period and potential time-dependent host factors during infection in the non-permissive host.

After incubation, plasmid library was extracted using ZR Plasmid Miniprep – Classic Kit (Zymo Research D4016) from control condition. Phage mixture was filtered using PES 0.22 µm filter, precipitated with the ammonium sulfate and Tween 80 method, DNase I treated, and purified using a DNA purification kit with the elution using TE buffer.

The relevant MOI was determined under the assumption that each member of the library would encounter a phage, and to avoid a scenario where multiple phages infect a single cell.

The selection experiment in *E. coli* K12 and B involved unpaired triplicates for both experimental and control conditions. However, we modified the experiment in *E. coli* O121 to a paired experimental design. This difference was considered in the subsequent analyses.

#### Plaque assay and efficiency of plating

The phage titer was quantified using double-overlay agar plaque assay. *E. coli* BW25113 was grown overnight in LB liquid media, back-diluted to OD_600_∼0.5 for plaque assay. We mixed 250 µl bacterial culture, 3.5 ml top agar (LB with 0.5% agar), and the relevant volume of phage lysate, vortexed, and plated on a pre-warmed LB agar plate. The solidified double-layer agar was incubated at 37°C for approximately 12 to 16 hours, unless otherwise stated. Efficiency of plating (EOP) for different bacterial host variants to wildtype *E. coli* was calculated as the ratio of T7 phage titer in the host variant (a gene knockdown) to the phage titer in permissive *E. coli* BW25113. The calculation considered the PFU within the range of 10-300, and the plaque assay was performed in triplicates. Statistical analysis was performed by Tukey’s honestly significant difference (HSD)(77) at the *p-*value of 0.05.

#### Bacteriophage purification and precipitation

Phage was purified post-infection by filtration through PES 0.22 µm syringe filter (30 mm diameter, Cell Treat 229747), and the phage lysate was stored at 4°C for the experiment.

We concentrated phage lysate using a two-step technique involving ammonium sulfate and Tween 80(78). Initially, 1 ml of filtered phage lysate was mixed with 100 µl 30% Tween 80 (final concentration ∼3%) and 350 mg ammonium sulfate (final concentration ∼35%w/v). After vigorous vortexing, the mixture was centrifuged at 3000xg for 20 minutes at 4°C. The top layer of the phage precipitate was scooped and resuspended in 100 µl TBS (20mM Tris, 150mM NaCl), then stored at 4°C.

#### Sample preparation, Sequencing, and Analysis

PCR preparation of samples for sequencing was conducted using KAPA HiFi (Roche KK2101) for a total of 24 cycles in a two-step PCR. The first PCR amplified the targeted sgRNA region with an addition of specific region for Illumina sequencing adaptor, and the second PCR attached sequencing indices to the two ends of the sequence (see Table S1). The sgRNA plasmid template amount varied; 5 ng of sgRNA library without selection and 10 µl of purified DNase-treated sgRNA plasmid after phage selection were used for the first PCR, and 1 µl PCR product without phage selection and 12 µl of purified PCR product were used for the second PCR. The template amount and volume depend on the type of samples and may need adjustment for an adequate amplification. PCR products were verified for the correct size using 1.2% agarose gel electrophoresis (0.6 g agarose, 50 ml 0.5x TAE with final concentrations of 20mM Tris-acetate, 0.5mM EDTA). PCR products were purified using E.Z.N.A® Cycle Pure Kits (Omega Bio-tek D6492-01) with a centrifugation protocol. DNA concentrating was assessed using a DNA concentrator (Zymo Research D4004) when the PCR product quantity was insufficient.

The targeted sgRNA was sequenced using a 2×150 bp paired-end Illumina sequencing system. Forward and reverse reads were paired using FLASH(79) (v1.2.11), filtered using VSEARCH(80) at e-value of 0.1, and aligned. The count of reads with each sgRNA was determined using MAGeCK(81). Data processing, analysis, and visualization were carried out using Python (3.8.0) and R in RStudio. Packages, including dplyr, tidyr, readxl (for data processing), genefilter, edgeR, limma, multtest, metap (for statistical analysis), ggrepel, ggplot2, gridExtra (for visualization), were used.

Significant changes in each sgRNA between the phage selection experiment and control were defined using edgeR(82) as employed in other CRISPR genetic screens(83–85), with a count per million (CPM) cutoffs that varied depending on the sequencing depth and the specific experiment. CPM of 6 was employed for the analysis of data in the pilot and experiment involving *E. coli* K12 and B. In the experiment with *E. coli* O121, a CPM threshold of 3 was applied due to a higher sequencing depth. This filtering strategy was specifically applied on the control condition. To ensure robustness across varying sequencing depth among hosts and experiments, the goal was to filter sgRNAs with a minimum of 3 reads for each sgRNA within each analysis and adjust the CPM accordingly. Briefly, edgeR estimates the statistical significance of the FC for each sgRNA before and after selection. The unpaired analysis was applied for data from *E. coli* K12 and B experiment, while the paired analysis was used for data obtained from *E. coli* O121 selection. The *p-*value was adjusted using False Discovery Rate (FDR)(86). Significance was determined by combining the *p-*values of each sgRNA targeted the same gene using Stouffer’s method(87), along with the log-transformed of mean FC of each gene and the distribution of the phage score caused by randomized sgRNAs. We employed two measurements of significance: one accounting for the variability of phage scores among sgRNAs targeting a gene, and the other assessing significance relative to a null distribution. The experiment was performed in triplicates and the scripts used are available on https://github.com/chitboonthav/CRISPRi_project.

Due to the unpaired design of the experiment in *E. coli* K12 and B, we assumed consistency across triplicates in the control. Experimental correlation was calculated by arbitrarily pairing each control with each selection result in pairwise combinations. The average log_2_FC was calculate for each experiment across all pairwise comparisons. The Pearson correlation coefficient was calculated to assess the correlation across replicates in experiments performed in all three strains (Supplementary Figure S4 and S5).

The mean and standard deviation of randomized sgRNAs were directly calculated for the phage score (log_2_FC) to establish a reference and background noise.

#### Individual gene knockdown by CRISPRi

Individual sgRNA oligos were purchased from IDT and resuspended to a concentration of 100µM ssDNA (see Table S1). For each sgRNA, two ssDNA with distinct overhangs relevant to the insertion site on the psgRNA01 vector were combined, forming dsDNA with a 5’-end overhang. Oligos were phosphorylated using T4 polynucleotide kinase (T4 PNK, NEB M0201S) in a reaction containing 1 µl of 100µM forward oligo, 1 µl of 100µM reverse oligo, 5 µl of 10x T4 ligase buffer (NEB B0202S), 1 µl of T4 PNK, and 42 µl of dH_2_O. The reaction was incubated at 37°C for 1 hour, followed by the addition of 2.5 µl of 1M NaCl. Subsequently, the mixture was incubated at 96°C for 6 minutes and 23°C for 30s with a ramp rate of 0.1°C/s, and then stored at −20°C. The phosphorylated dsDNA oligo was 10-fold diluted for Golden Gate Assembly. This reaction included mixing 75 ng of linearized psgRNA01 backbone, 1 µl of diluted dsDNA oligo, 2 µl of T4 ligase buffer, 1 µl of Golden Gate Enzyme Mix (BsaI-HFv2, NEB M2616AA), and adding dH_2_O to reach a total volume of 20 µl. The reaction was incubated at 37°C for 1 hour, heat inactivated at 60°C for 5 minutes, dialyzed for 1 hour, transformed into a host with integrated *dCas9* genome, and selected on Crb-Km agar plate. Transformants were verified through colony PCR and Sanger sequencing (Functional Biosciences Madison, Wisconsin).

An EOP assay, as described earlier, was performed to quantify the gene knockdown effect on T7 plaquing efficiency compared to wildtype *E. coli* O121 and BW25113. CRISPRi was induced overnight (1mM IPTG), and 200 µl of the overnight bacterial culture was used for the plaque assay. The number of plaques was counted after 4-6 and 16 hours of incubation at 37°C.

#### Querying *E. coli* genes in a curated IMG/VR v4 database

The IMG/VR-v4 database was filtered for high-quality viruses by querying the “MIUViG quality” field. These high-quality viruses were dereplicated by selecting a representative sequence from each viral operational taxonomic unit (vOTU) with the highest estimated genome completion and lowest estimated genome contamination. All *E. coli* genes from strains BL21 (RefSeq: NZ_CP053601) and BW25113 (RefSeq: NZ_CP009273) were then queried against the filtered IMG/VR-v4 database(64) using Diamond(88) (v2.1.0) with settings “--evalue 1e-5 --more-sensitive -k0 --id 50”. Hits with coverage ≥50% to the viral protein were considered true hits.

To eliminate prophage genes in the *E. coli* genomes, each *strain’s E.* coli genes were annotated using the PHROGs(89) database (downloaded 2022-01-17) via mmseqs(90) (v13.45111), following the settings established in the PHROGs documentation. Any *E. coli* gene identified by PHROGs database with an e-value ≤ 1e-3 and at least 37% coverage against the viral protein profile was considered a true viral protein. Subsequently, all genes identified through PHROGs annotation were categorized as viral or bacterial based on their abundance in the viral genomes calculated earlier. The elbow point of a percentile plot (percentile rank vs percentile threshold value, see Supplementary Figure S13) of the PHROGs-annotated gene counts in viral genomes was used as the threshold to partition these genes into viral and bacterial genes. The elbow point of this plot was calculated using the kneed (v0.8.5) Python(91) package. Genes falling below this threshold count in viral genomes and any *E. coli* gene not identifiable by annotation with PHROGs were considered true *E. coli* genes.

#### Statistical significance of *E. coli* gene counts in viral genomes

The statistical significance of *E. coli* gene counts found in viral genomes was assessed using a Mann-Whitney U (MWU) test(92) implemented in the SciPy(93) (v1.10.1) Python package. The null hypothesis posits that the distribution of gene counts in viral genomes does not differ for genes impacting phage score compared to those that do not. For each combination of *E. coli* strains (K-12, B) and phage score outcomes (increased/decreased), two paired sets of gene counts were sampled from the set of genes that did not change phage score and from the corresponding set of genes that influenced phage score during CRISPRi knockdown. A sample size equal to 60% of the size of the gene set corresponding to the score category was taken, rounded to the nearest integer. This process was repeated 10,000 times, and the average number of viral homologs was stored for each iteration. The distribution for a given score category was compared to that of the paired distribution for the set of genes not impacting phage score in the given strain using the MWU test with a significance cutoff of 0.05.

#### Protein structure prediction

We predicted the protein structure using ColabFold(55, 56) (v1.5.3, AlphaFold2 using MMseqs2). Briefly, we followed the documentation and provided the amino acid sequences of the proteins of interest into the program. The quality of predicted protein structures was assessed through Multiple Sequence Alignment (MSA) sequence coverage plots and the Predicted Aligned Error (PAE) (Supplementary Figure S12). We selected the top-ranked structures. All protein structures were visualized and examined using PyMOL. We searched and compared the protein structure using MPI Bioinformatics Toolkit(94, 95).

All supplementary data is available in Table S8.

## Author contribution

CC, PH and SR conceptualized and designed the study. CC constructed plasmids and the bacterial sgRNA library, conducted the experiment, curated the data, performed data analyses, designed the figures, and wrote the manuscript. CM and KA carried out bioinformatic analysis of viral genomes. JMP assisted with the CRISPRi screen. All authors reviewed and edited the manuscript, provided critical feedback, and contributed to discussions on the manuscript.

## Acknowledgment

CC was supported by the Anandamahidol Foundation Scholarship, Thailand. SR was supported by the National Institute of Allergy and Infectious Disease under award R21AI156785 and National Science Foundation CAREER Award. The work of CM and KA was partly supported by the National Institute of General Medical Sciences (NIGMS)/NIH to KA (R35GM143024). CM was supported by a US National Science Foundation Graduate Research Fellowship. JMP was supported by the National Institute of General Medical Sciences of the National Institutes of Health under award number R35GM150487. We thank Agilent Technologies for providing SurePrint Oligonucleotide libraries and Laura Whitman for oligo synthesis support. The content is solely the responsibility of the authors and does not necessarily represent the official views of the National Institutes of Health or the National Science Foundation.

## Notes

### Competing Interest Statement

The authors have declared no competing interest.

