## Supplementary_Figures for "Systematic genome-wide discovery of host factors governing bacteriophage infectivity"

**Supplementary Figure S1** Development of PHAGEPACK, integrating CRISPRi and phage packaging strategies
**Supplementary Figure S2** The coverage of sgRNA in the library construction targeting *E. coli* K12 and B strain
**Supplementary Figure S3** Randomized sgRNA showed no effect on phage
**Supplementary Figure S4** The correlation of replicates in *E. coli* K12
**Supplementary Figure S5** The correlation of replicates in *E. coli* B
**Supplementary Figure S6** The distribution of sgRNA library targeting *E. coli* O121 in 10G
**Supplementary Figure S7** Randomized sgRNA exhibited no change of phage score in *E. coli* O121
**Supplementary Figure S8** The correlation of replicates in *E. coli* O121 at 45-minutes selection
**Supplementary Figure S9** Phage scores in *E. coli* O121 under a 2-hour selection
**Supplementary Figure S10** The correlation of replicates in *E. coli* O121 at 2-hours selection
**Supplementary Figure S11** Three hypothetical proteins predicted by ColabFold (AlphaFold2)
**Supplementary Figure S12** ColabFold (AlphaFold2) parameter for protein predictions
**Supplementary Figure S13** Viral genome prediction
**Supplementary Figure S14** qPCR primer efficiency

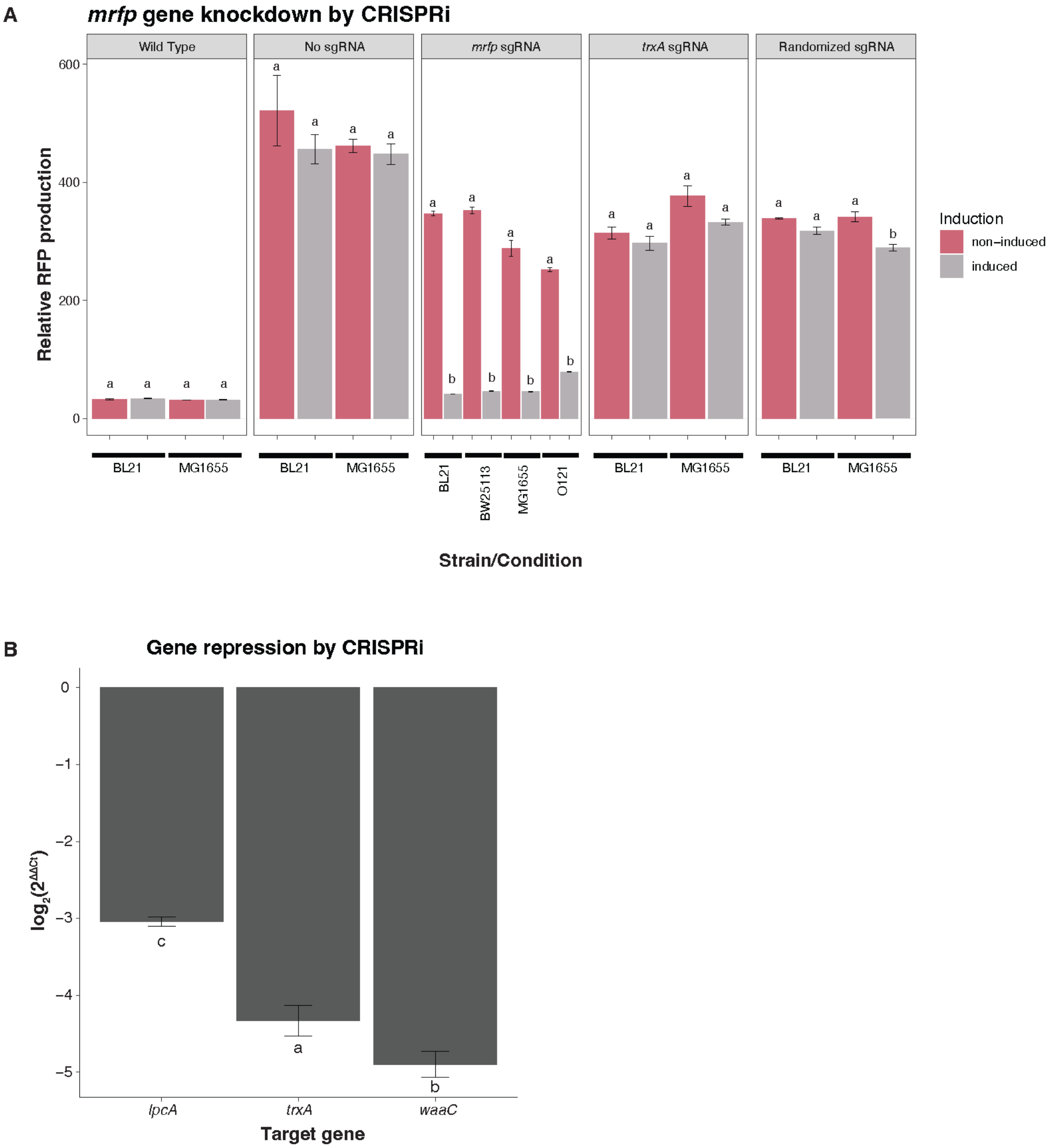


#### Supplementary Figure S1 Development of PHAGEPACK, integrating CRISPRi and phage packaging strategies

(A) The *mrfp* gene was genome integrated into the target bacterial hosts considered in this study. The validation of gene knockdown was confirmed through the suppression of *mrfp* gene, with a subsequent assessment of mRFP production. The results, reported as relative RFP production, were examined across various strains and conditions. Colors represent the induction conditions, distinguishing between non-induced (pink) and induced (grey) states. The conditions were categorized into wildtype and hosts containing *dCas9* and *mrfp* genes. Additionally, the differences in sgRNA present in the cell were considered, encompassing no sgRNA, *mrfp*-target, *trxA*-targeted*,* and randomized sgRNA. (B) The validation of targeted gene knockdown was evaluated through RT-qPCR of the transcripts when *lpcA*, *trxA*, and *waaC* were specifically targeted. All data are reported as the mean±SD across biological replicates. Host receptors for bacteriophage adsorption(77, 96). Statistical analysis was performed by Student's *t*-test and Tukey's HSD test, respectively at the *p*-value of 0.05(77, 96).

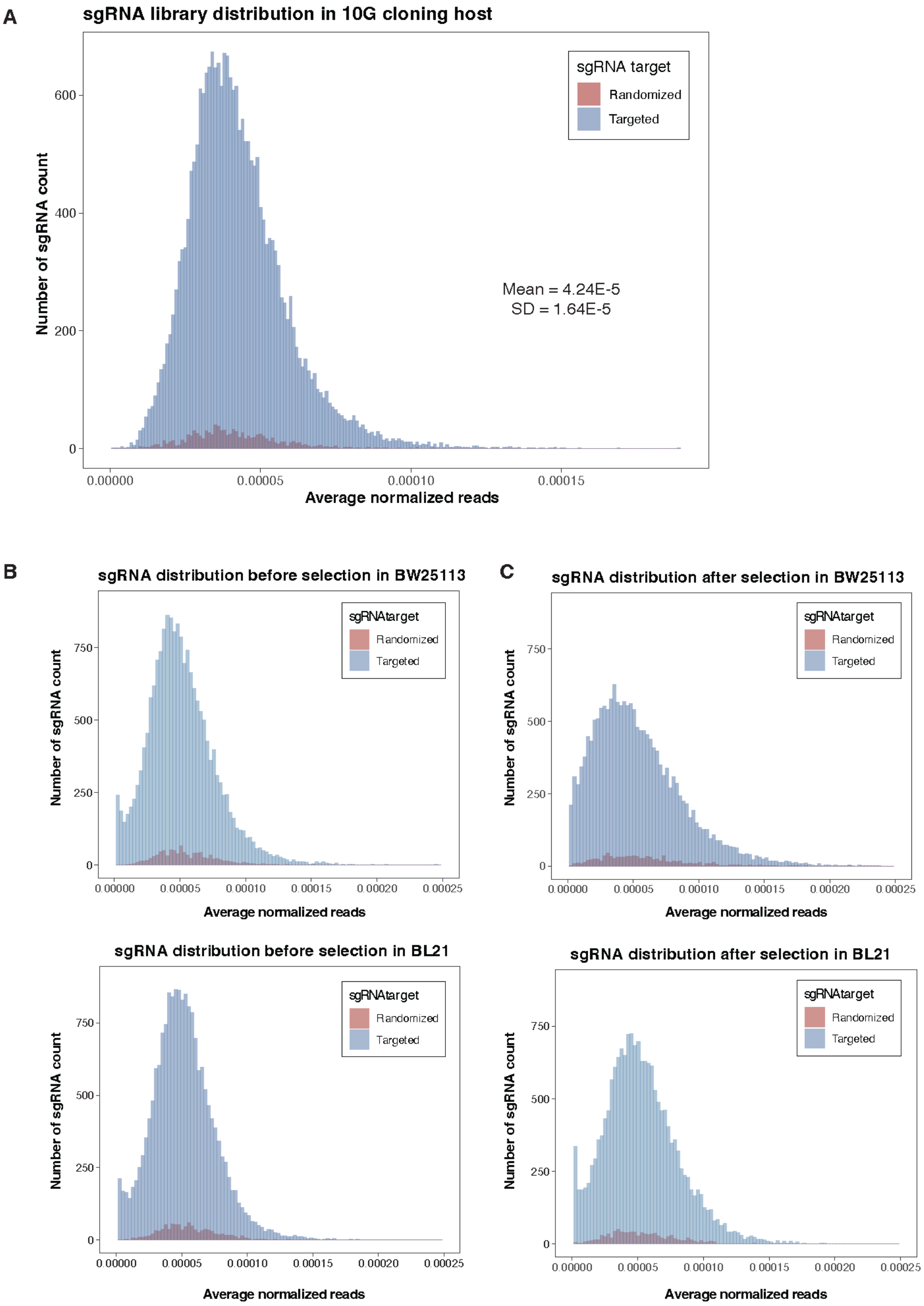
A
sgRNA library distribution in 10G cloning host
sgRNA target
Randomized
Targeted
Mean = 4.24E-5
SD = 1.64E-5
Number of sgRNA count
0
200
400
600
0.00000
0.00005
0.00010
0.00015
Average normalized reads
B
sgRNA distribution before selection in BW25113
C
sgRNA distribution after selection in BW25113
sgRNAtarget
Randomized
Targeted
Number of sgRNA count
0
250
500
750
0.00000
0.00005
0.00010
0.00015
0.00020
0.00025
Average normalized reads
sgRNA distribution before selection in BL21
sgRNA distribution after selection in BL21

#### Supplementary Figure S2 The coverage of sgRNA in the library construction targeting *E. coli* K12 and B strains

(A) The sgRNA library was transformed into wildtype *E. coli* 10G cloning strain to ensure an unbiased library construction, resulting in complete coverage of all 19,471 sgRNAs, which exhibited a normal distribution. (B) The sgRNA library in targeted hosts, K12 (BW25113; Top) and B (BL21; Bottom), exhibited a normal distribution prior to phage selection. (C) The distribution of sgRNA library in K12 (BW25113; Top) and B (BL21; Bottom) following phage selection.

**A**

**Distribution of $log_2FC$ of randomized sgRNA in BW25113 selection (CPM6)**

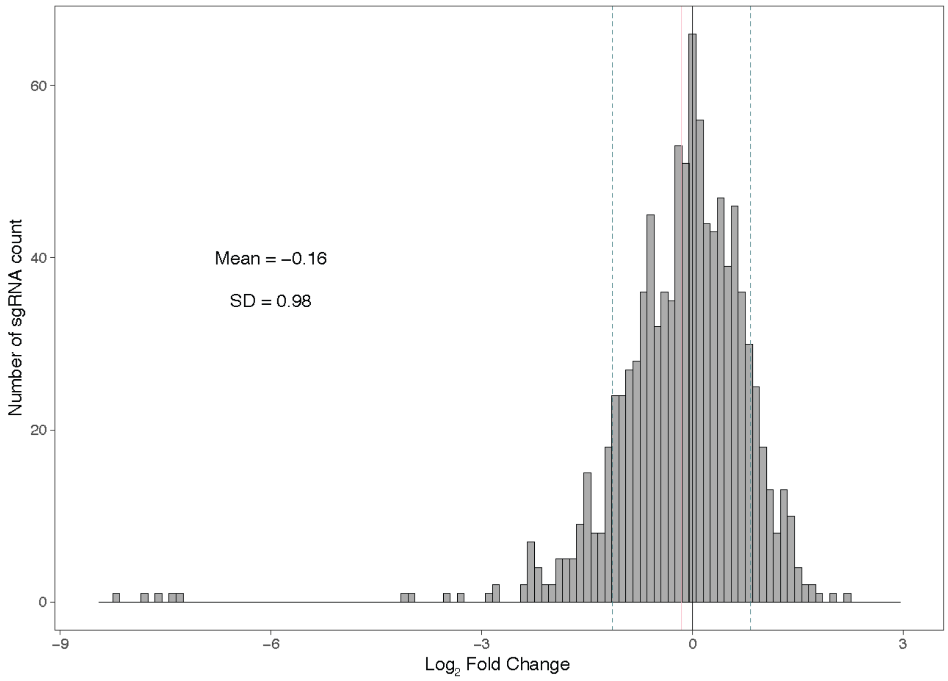


**B**

**Distribution of $log_2FC$ of randomized sgRNA in BL21 selection (CPM6)**

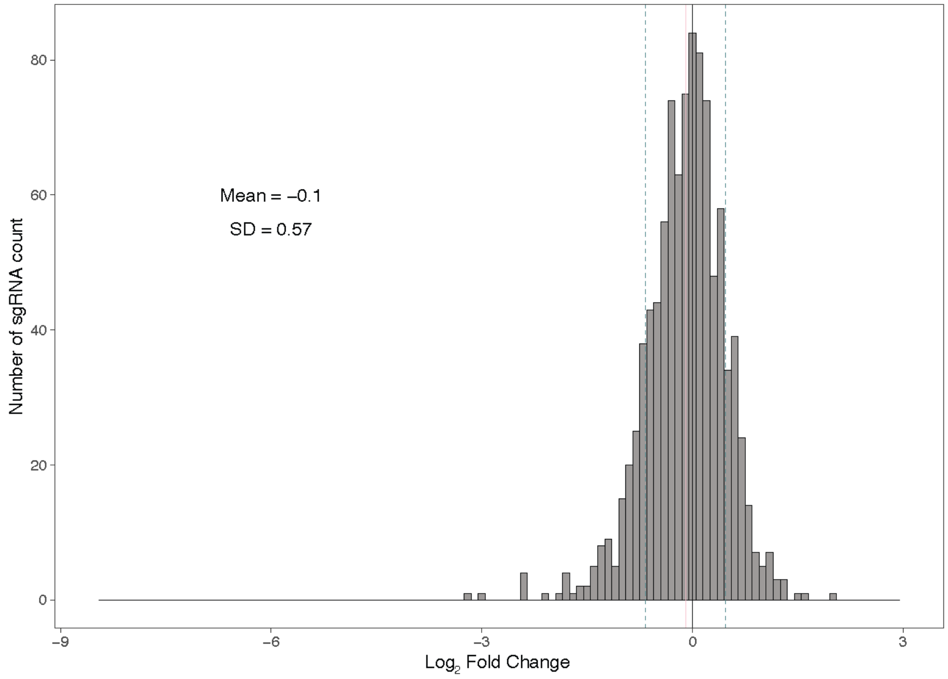

#### Supplementary Figure S3 Randomized sgRNA shows no effect on phage

The distribution of phage functional scores ($log_2FC$) resulting from randomized sgRNAs in *E. coli* K12 (BW25113; A) and B (BL21; B). Vertical solid pink and blue dashed lines indicate the mean and the first standard deviation.

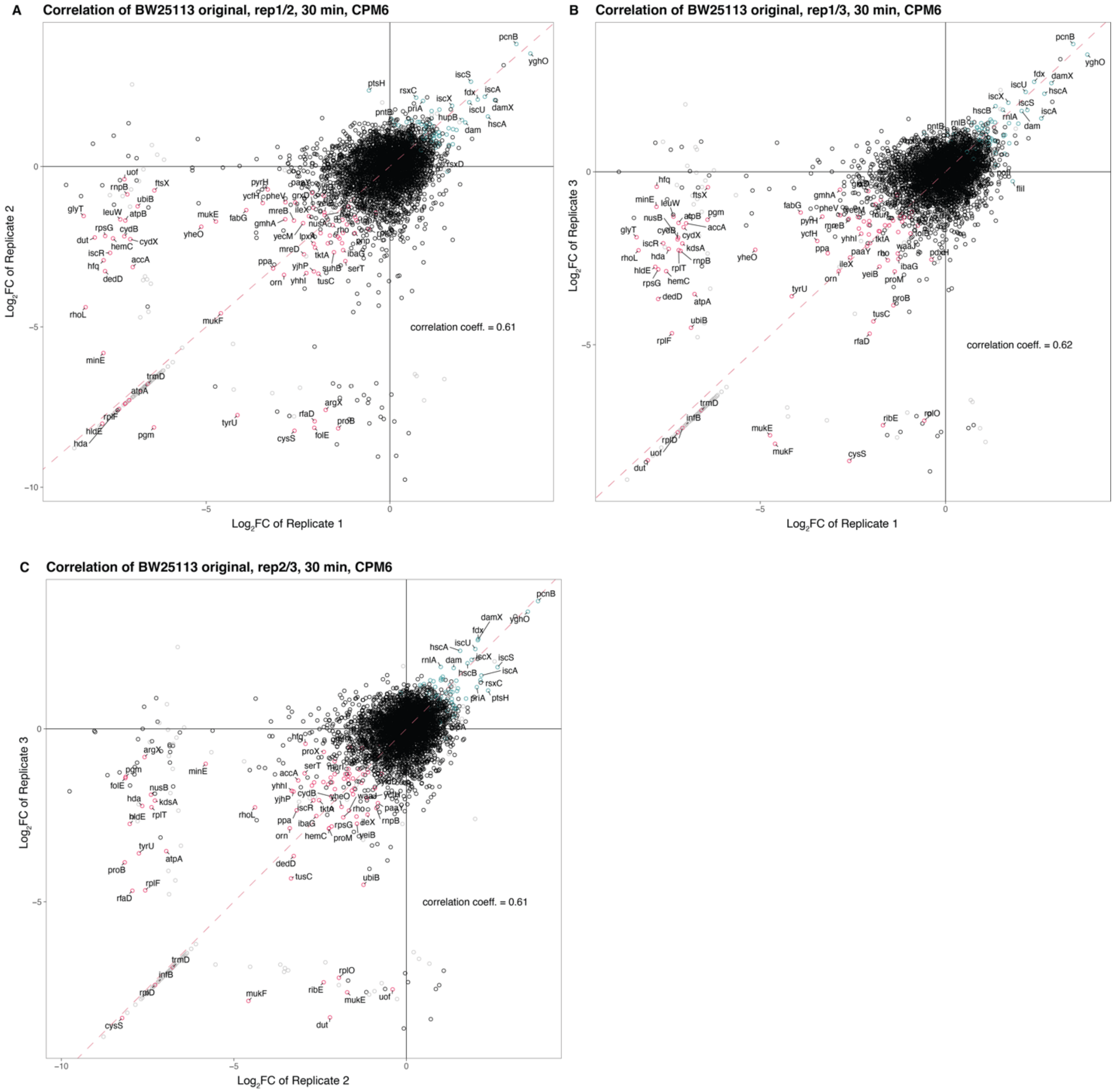


### Supplementary Figure S4 The correlation of replicates in *E. coli* K12

The correlation of phage scores across replication in the phage selection experiment is illustrated among (A) replicates 1 and 2, (B) replicates 1 and 3, and (C) replicates 2 and 3. Each dot represents an individual bacterial gene, color-coded based on the significance of its impact on phage scores – increasing, decreasing, or neutral in green, red, and black, respectively. Genes with knockdown that did not pass the filtering criteria are shown in grey. The correlation coefficient was calculated using Pearson's correlation. Diagonal dashed line indicates x is equal to y.

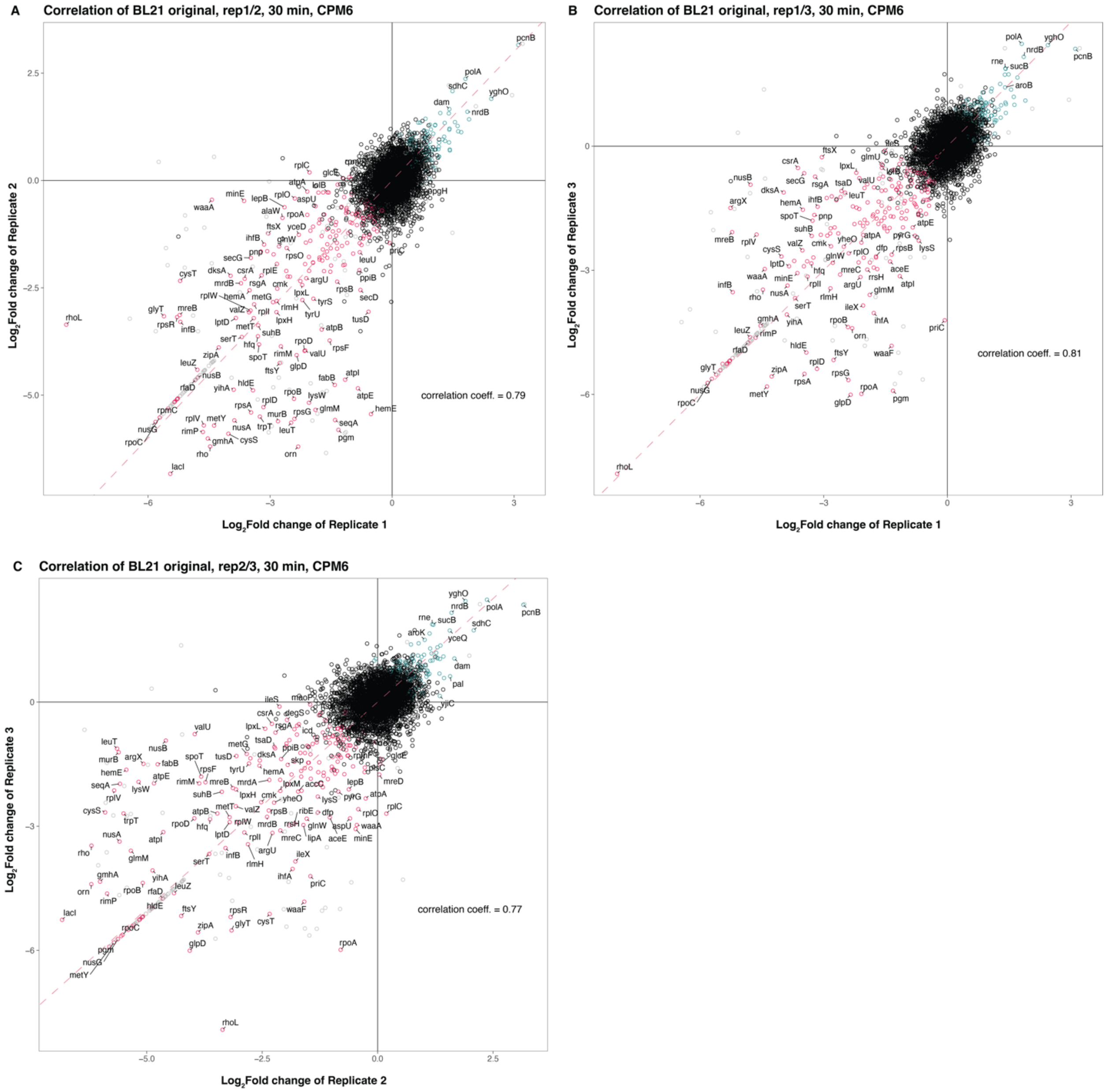


### Supplementary Figure S5 The correlation of replicates in *E. coli* B.

The correlation of phage scores across replication in the phage selection experiment is illustrated among (A) replicates 1 and 2, (B) replicates 1 and 3, and (C) replicates 2 and 3. Each dot represents an individual bacterial gene, color-coded based on the significance of its impact on phage scores – increasing, decreasing, or neutral in green, red, and black, respectively. Genes with knockdown that did not pass the filtering criteria are shown in grey. The correlation coefficient was calculated using Pearson's correlation. Diagonal dashed line indicates x is equal to y.

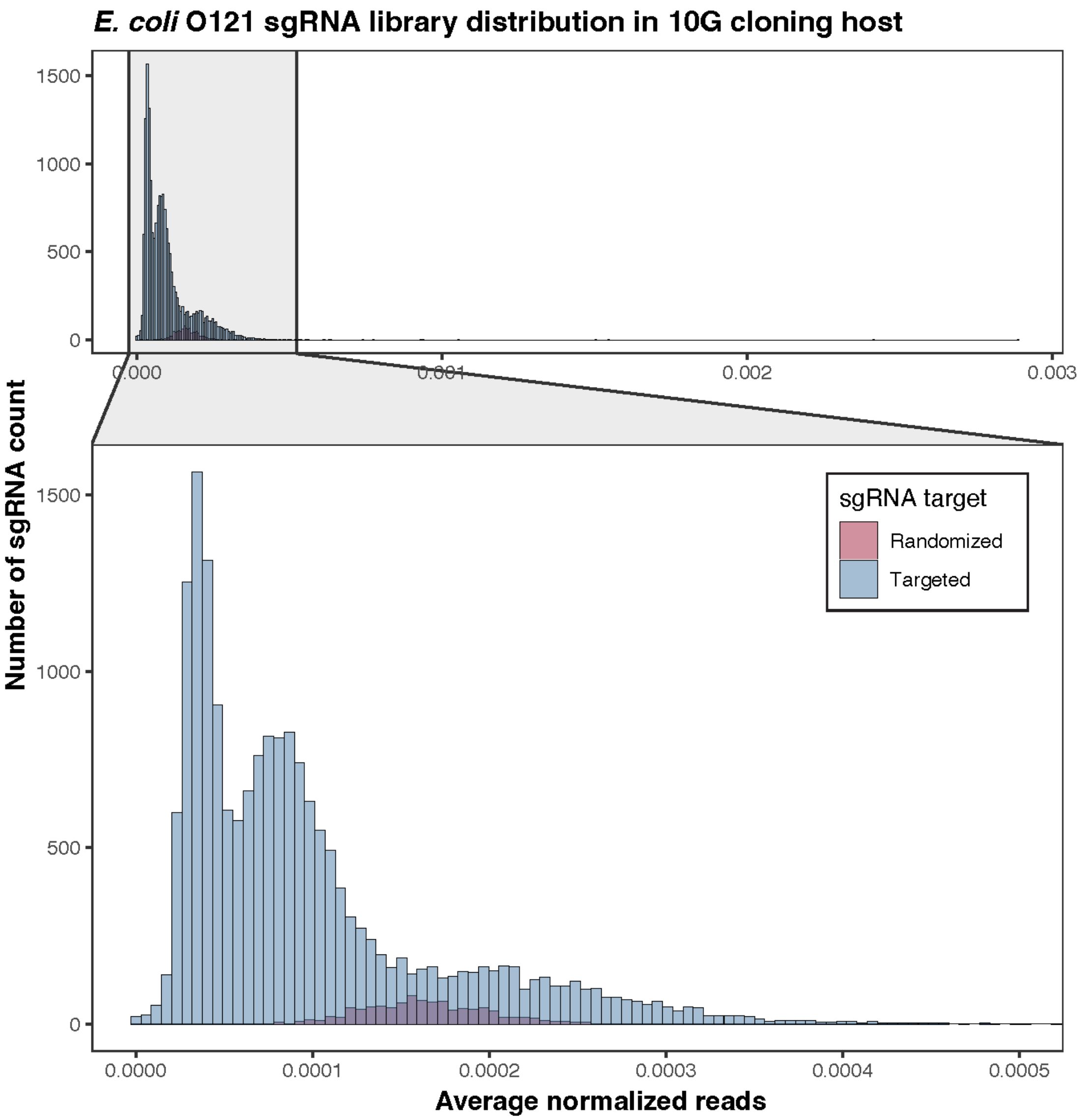


#### Supplementary Figure S6 The distribution of sgRNA library targeting *E. coli* O121 in *E. coli* 10G

The sgRNA library targeting *E. coli* O121 was transformed into wildtype *E. coli* 10G cloning strain, colored by type of sgRNA as Targeted (blue) and Randomized (red).

A

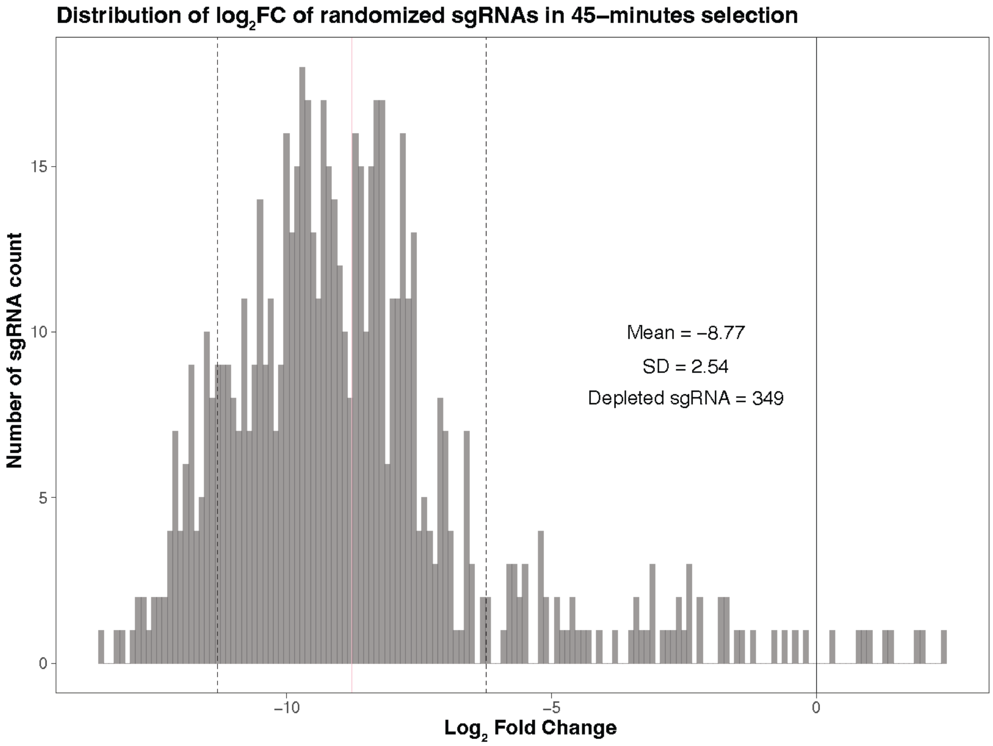
Distribution of $log_2$FC of randomized sgRNAs in 45–minutes selection
Mean = −8.77
SD = 2.54
Depleted sgRNA = 349
Number of sgRNA count
15
10
5
0
−10
−5
0
$Log_2$ Fold Change

B

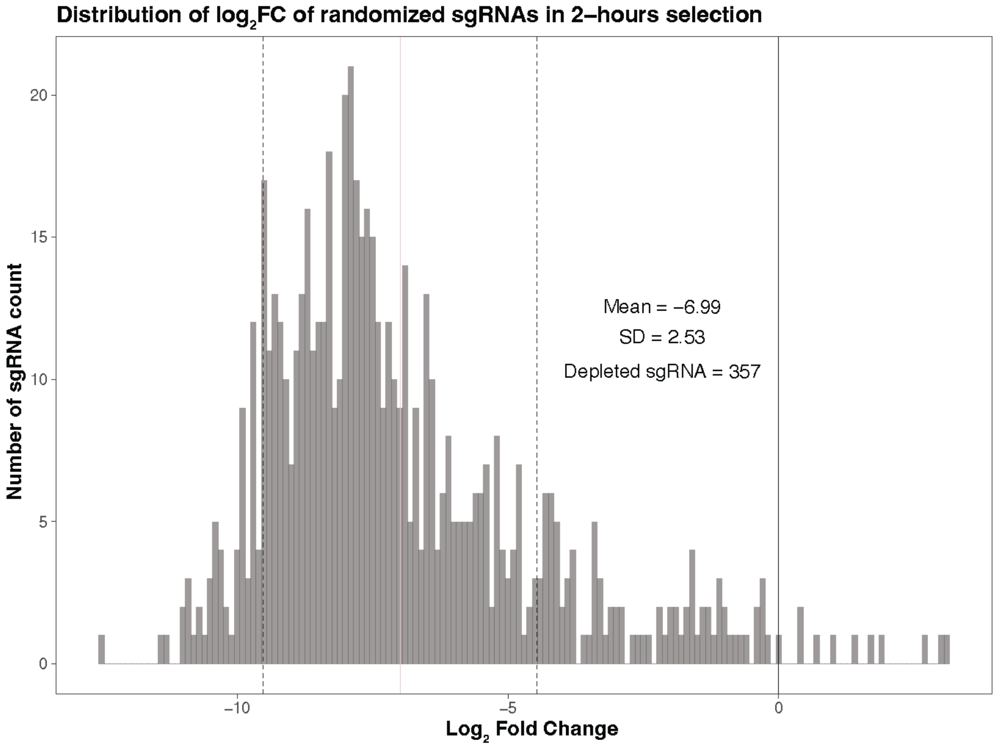
Distribution of $log_2$FC of randomized sgRNAs in 2–hours selection
Mean = −6.99
SD = 2.53
Depleted sgRNA = 357
Number of sgRNA count
20
15
10
5
0
−10
−5
0
$Log_2$ Fold Change

#### Supplementary Figure S7 Randomized sgRNA exhibited no change of phage score in *E. coli* O121

The distribution of phage functional scores resulting from randomized sgRNAs in *E. coli* O121 after a phage selection at (A) 45 minutes and (B) 2 hours. Vertical solid and dashed lines indicate the mean and the standard deviations (SD).

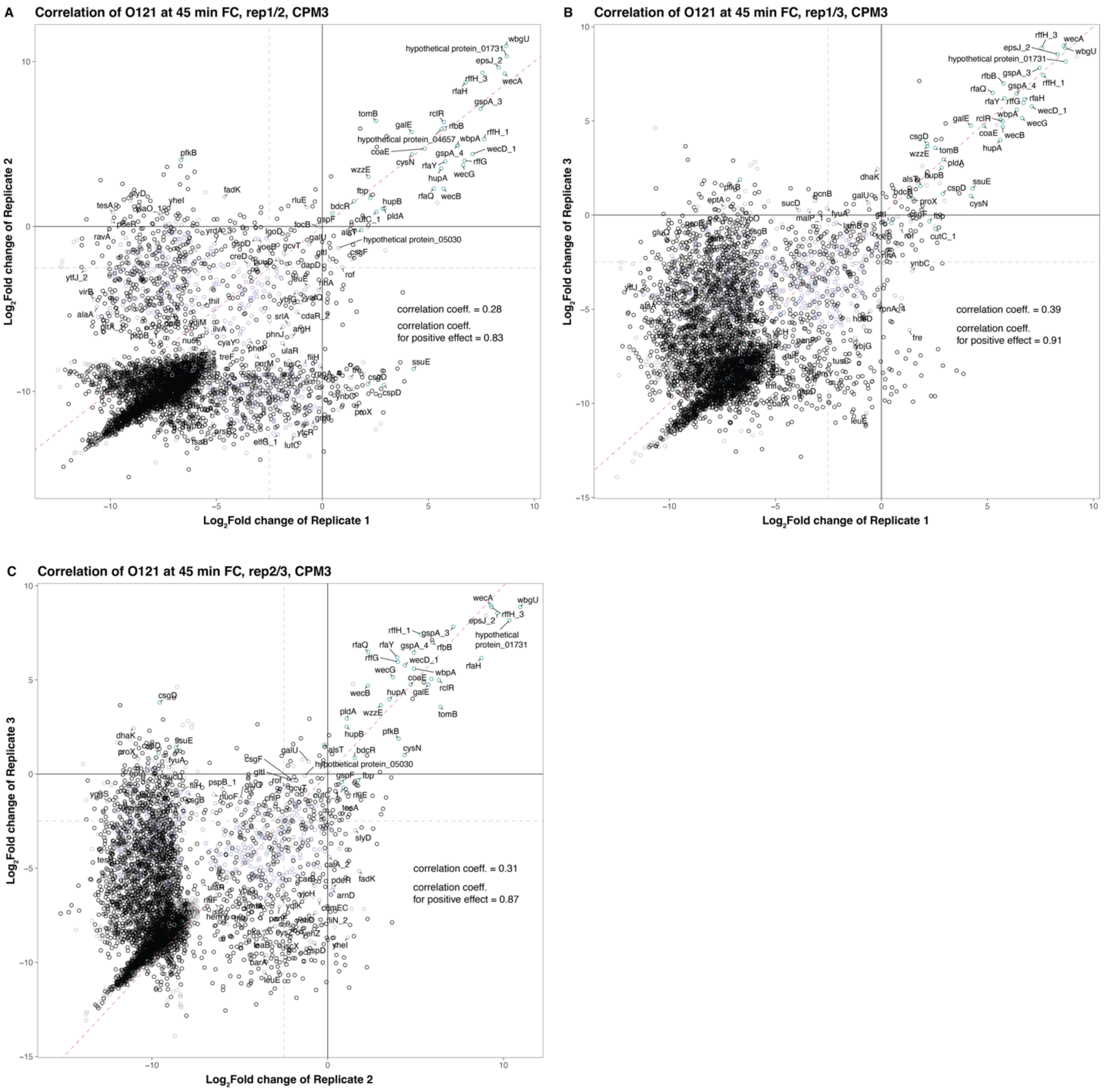


### Supplementary Figure S8 The correlation of replicates in *E. coli* O121 at 45-minutes selection

The correlation of phage scores across replicates in the phage selection experiment is illustrated between (A) replicates 1 and 2, (B) replicates 1 and 3, and (C) replicates 2 and 3. Each dot represents individual bacterial gene, color-coded based on the impact of the gene knockdown on the phage scores – increasing, detectable but decreasing, and neutral in green, purple, and black. Grey indicates the gene knockdowns that did not pass the filtering criteria. Diagonal dashed line indicates x is equal to y. Horizontal and vertical dashed lines reference a $log_2FC$ of -2.5. The correlation coefficient was calculated using Pearson's correlation. The correlation coefficients were calculated for the entire set of genes (correlation coeff.), and specifically for genes with a phage score larger than -2.5 (correlation coeff. for positive effect).

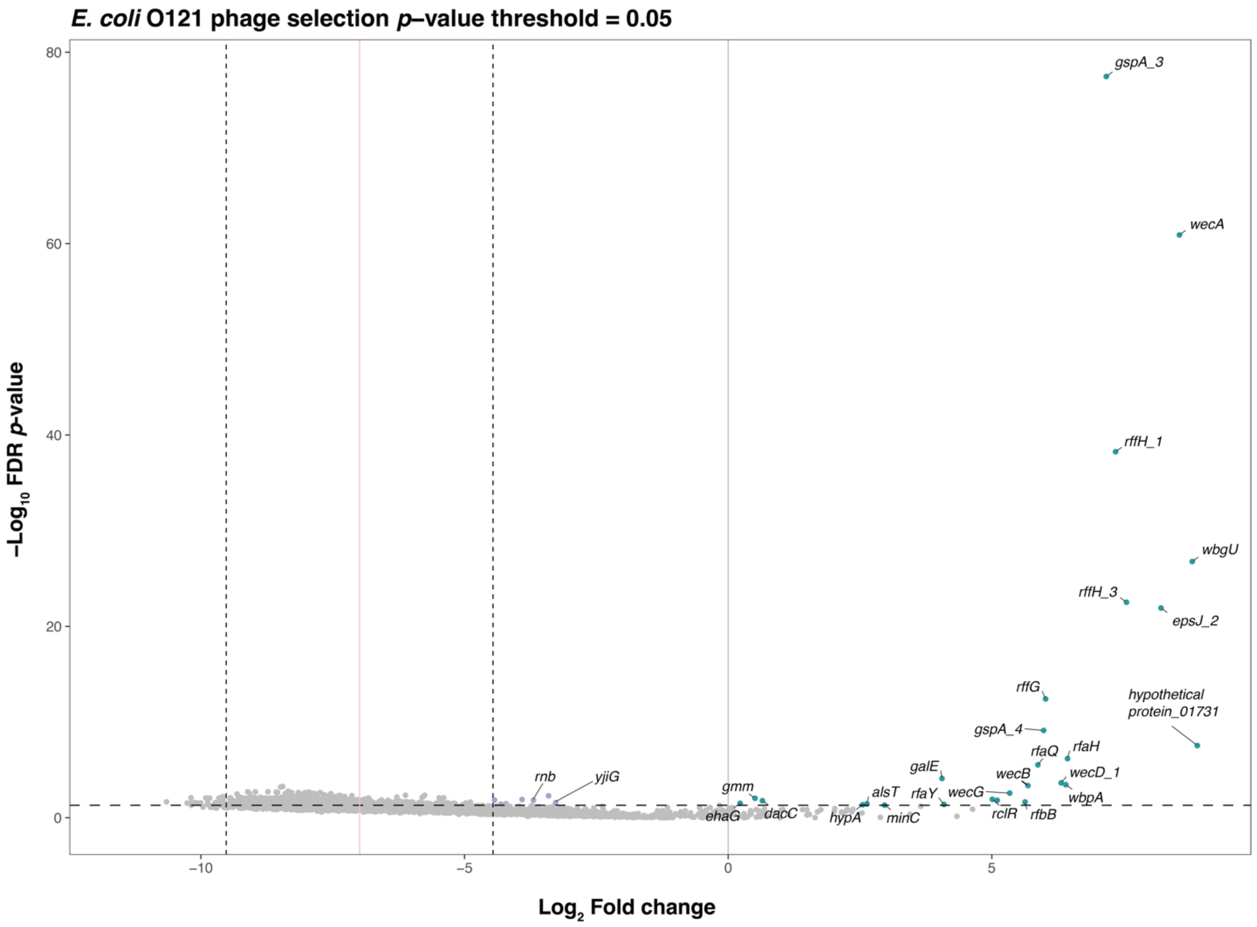


### Supplementary Figure S9 Phage scores in *E. coli* O121 under a 2-hour selection.

Phage scores in *E. coli* O121 after 2-hours of phage selection. Each dot is colored in green, purple, or grey, representing the knockdown of individual genes that significantly increased, detectable but smaller than 0, or showed no impact (smaller than -2.5) on the phage score. Vertical lines, pink solid, dashed black and grey lines, represent the mean, standard deviations, and zero.

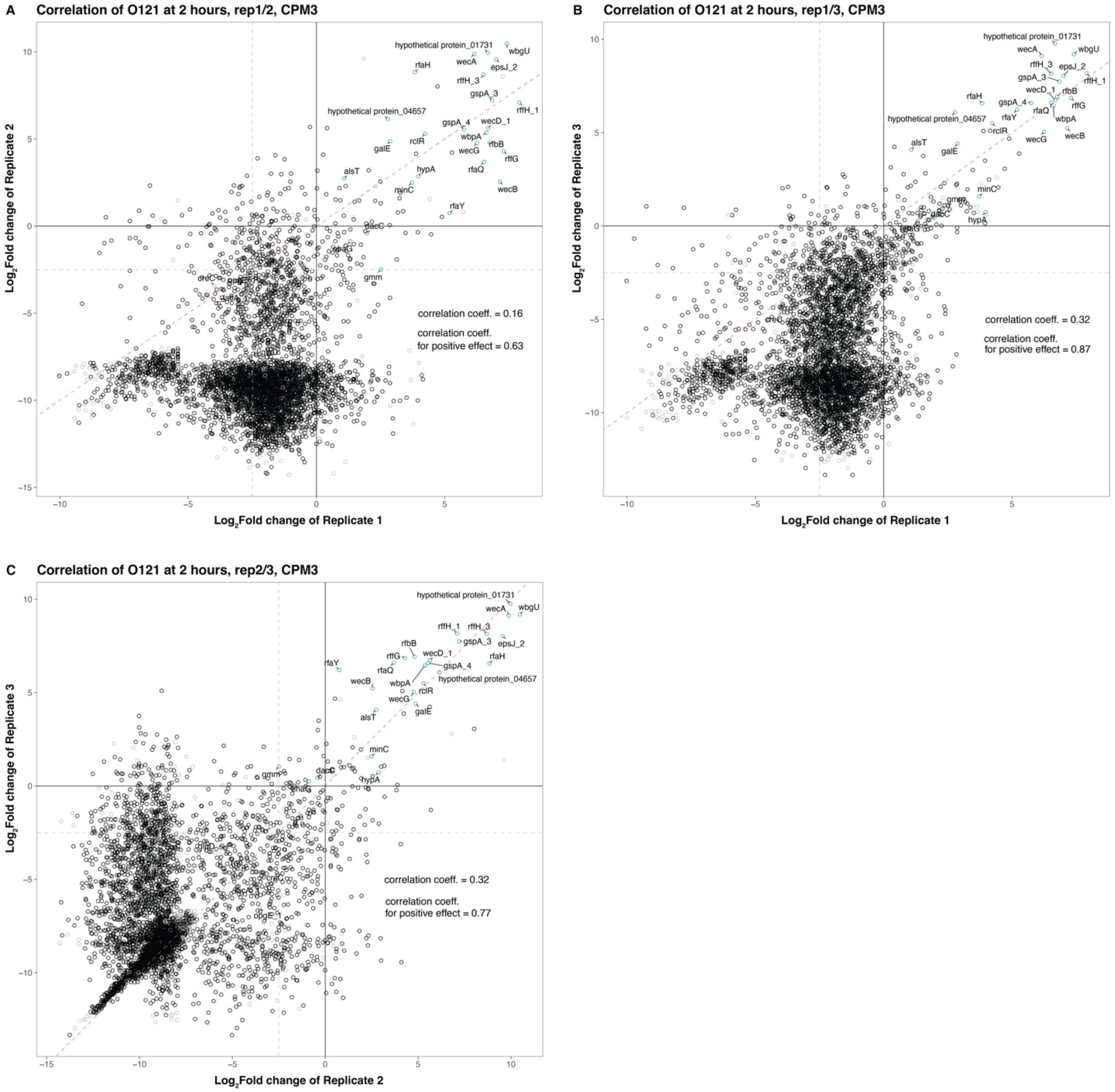


### Supplementary Figure S10 The correlation of replicates in *E. coli* O121 at 2-hours selection.

The correlation of phage scores across replicates in the phage selection experiment is illustrated between (A) replicates 1 and 2, (B) replicates 1 and 3, and (C) replicates 2 and 3. Each dot represents individual bacterial gene, color-coded based on the impact of the gene knockdown on the phage scores – increasing, detectable but decreasing, and neutral in green, purple, and black. Grey indicates the gene knockdowns that did not pass the filtering criteria. Diagonal dashed line indicates x is equal to y. Diagonal dashed line indicates x is equal to y. Horizontal and vertical dashed lines reference a $\log_2$FC of -2.5. The correlation coefficient was calculated using Pearson's correlation. The correlation coefficients were calculated for the entire set of genes (correlation coeff.), and specifically for genes with a phage score larger than -2.5 (correlation coeff. for positive effect).

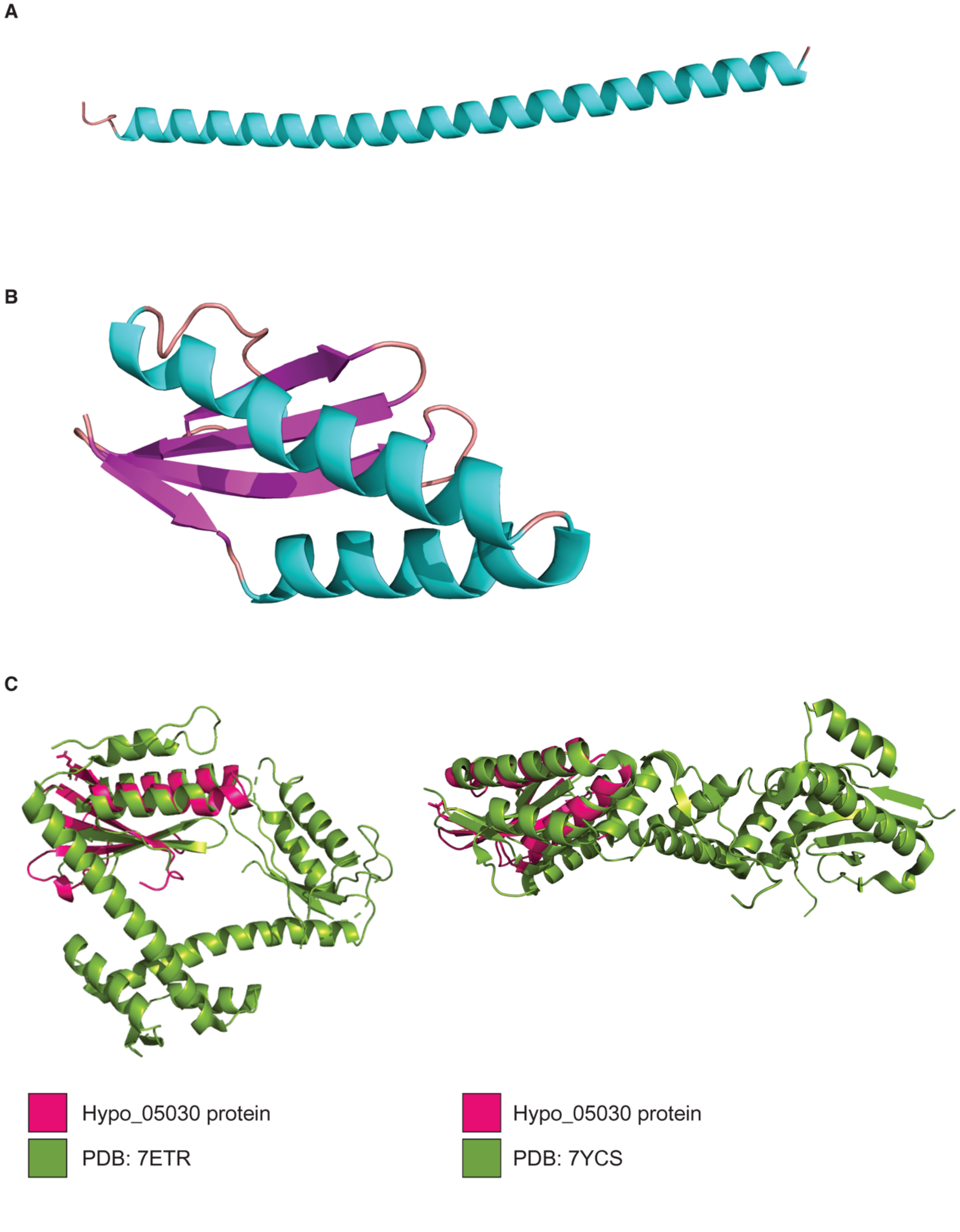
A
B
C
Hypo_05030 protein
PDB: 7ETR
Hypo_05030 protein
PDB: 7YCS

#### Supplementary Figure S11 Three hypothetical proteins predicted by ColabFold (AlphaFold2)

Predicted structures ranked 1 of (A) hypothetical proteins 01731 and (B) 05030 by ColabFold. The structures are color-coded to represent different structural elements: blue for $\alpha$-helix, magenta for β-sheet, and light pink for a loop. The structural alignments of hypothetical protein 05030 (pink) and ParE toxin at 2.296Å (PDB:7ETR, left), and ParD antitoxin at 4.522Å (PDB:7YCS, right).

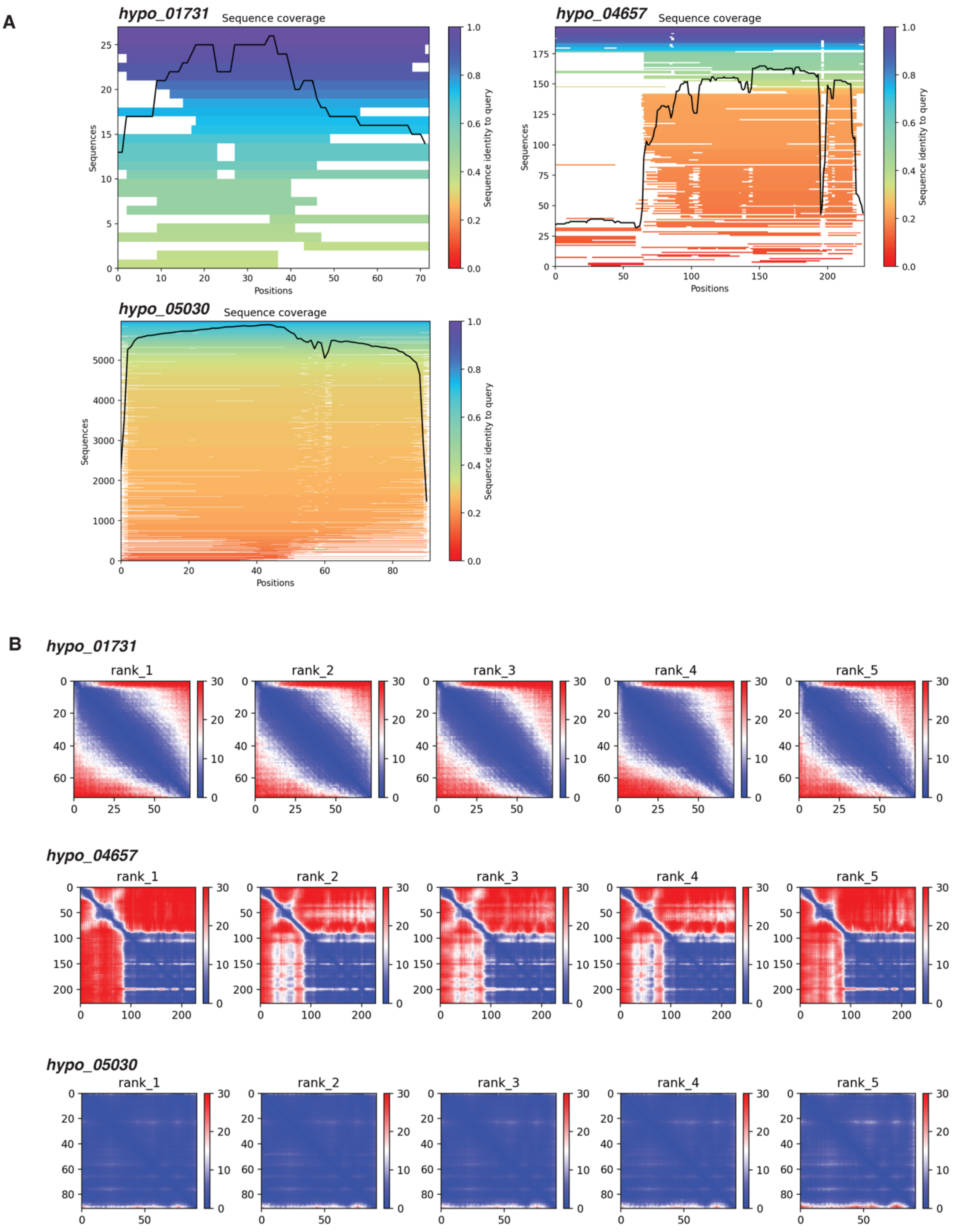
A
hypo_01731
Sequence coverage
Sequences
Positions
Sequence identity to query
hypo_04657
Sequence coverage
Sequences
Positions
Sequence identity to query
hypo_05030
Sequence coverage
Sequences
Positions
Sequence identity to query
B
hypo_01731
rank_1
rank_2
rank_3
rank_4
rank_5
hypo_04657
rank_1
rank_2
rank_3
rank_4
rank_5
hypo_05030
rank_1
rank_2
rank_3
rank_4
rank_5

#### Supplementary Figure S12 ColabFold (AlphaFold2) parameter for protein predictions

(A) Multiple Sequence Alignment (MSA) sequence coverage plots obtained from protein predictions of hypothetical proteins (Hypo_01731, Hypo_04657, Hypo_05030) by ColabFold. Each bar in the plots represents the number of aligned sequences at each position. The color intensity of the bars indicates the similarity of the sequences to the query, with bluer bars representing higher similarity. The black line indicates the number of the sequence matches to the query at each position. (B) The Predicted Aligned Error (PAE) graphically represents the confidence of residue-residue interactions, ranging from high at 0 (blue) to low at 30 (red). PAE was reported for top five predicted protein structures ranked from 1 to 5 (as rank_1 to rank_5).

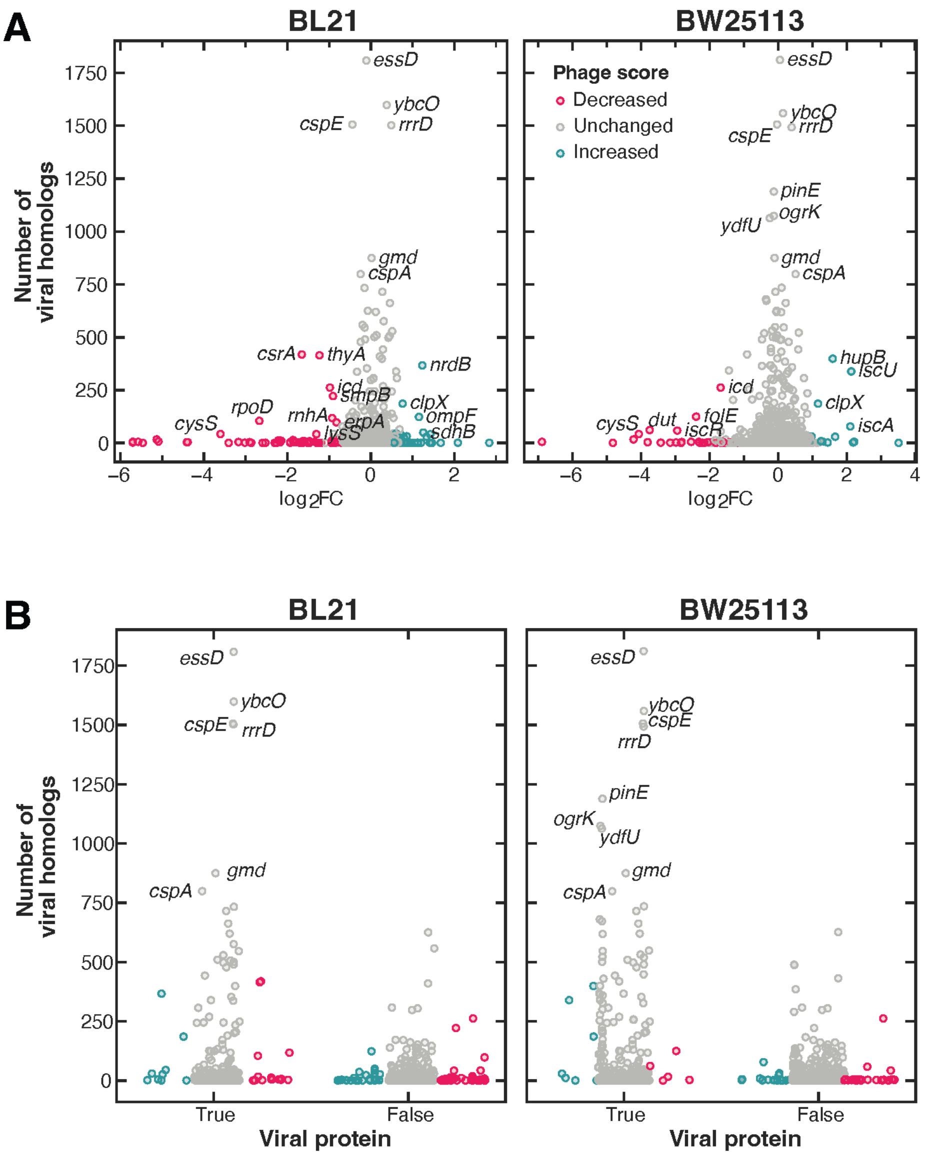


### Supplementary Figure S13 Viral genome prediction

(A) Experimental phage scores, represented in $\log_2$ fold change for individual genes, were compared to the number of viral homologs corresponding to the *E. coli* BW25113 (right) and BL21 (left) genes. Genes were color-coded as red (decreased), grey (unchanged), and green (increased), based on their phage scores in PHAGEPACK screen. (B) All *E. coli* genes were annotated using virus-specific gene annotation methods to identify potential viral genes in the *E. coli* genome. Genes specific to viruses or relevant to prophages were labeled as True, while *E. coli* genes unrelated to viruses were labeled as False. Each dot in this figure, representing individual genes, was color-coded: red for a significant decrease, green for a significant increase, and grey for a neutral effect due to gene knockdown on phage score.

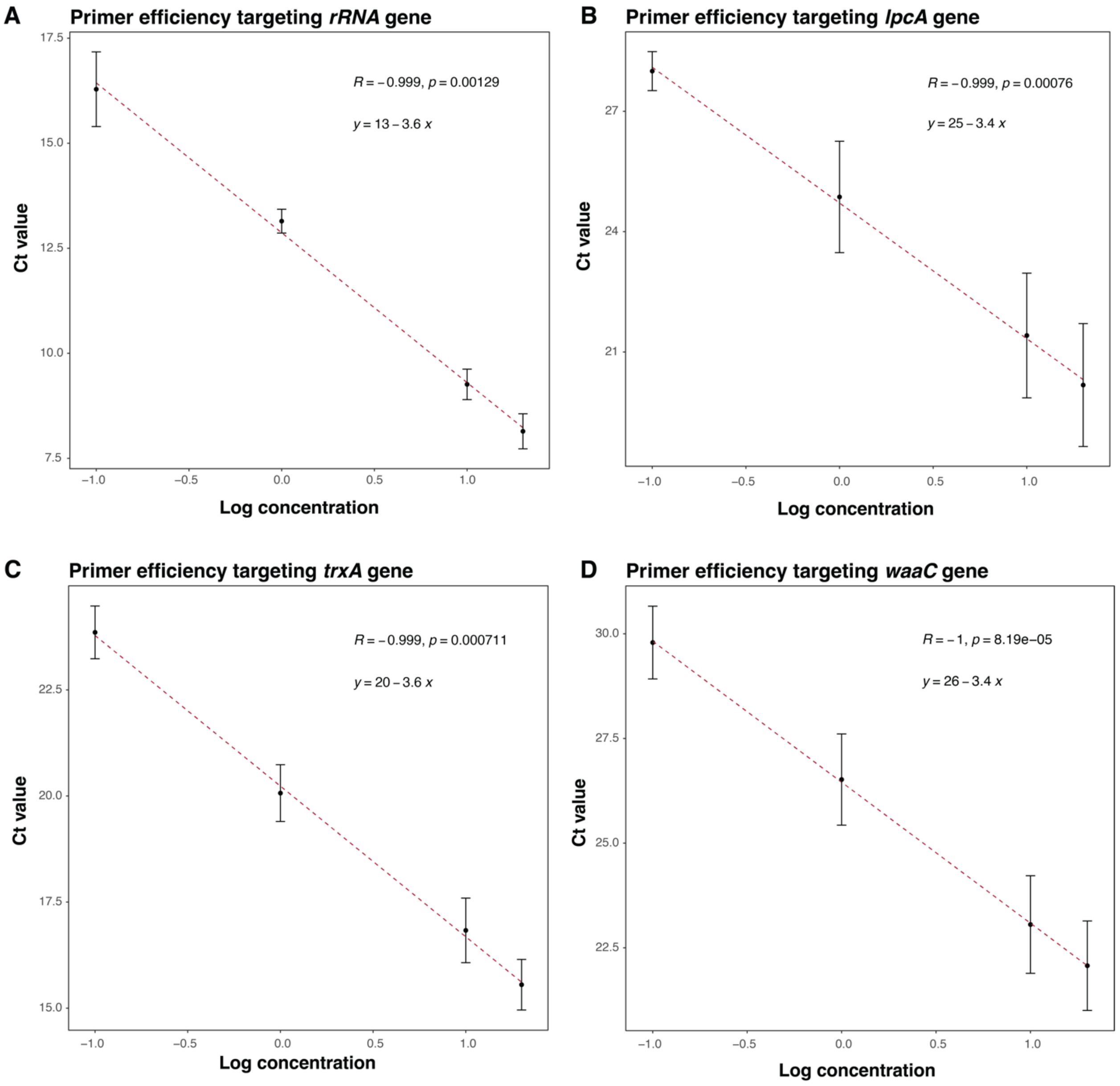


#### Supplementary Figure S14 qPCR primer efficiency

The efficiency of primers for RT-qPCR quantification of gene transcripts, including (A) *rRNA*, (B) *lpcA*, (C) *trxA*, and (D) *waaC*, was assessed.

### Supplementary Tables

**Table S1:** Sequences of primers, plasmids, and oligos
**Table S2:** Sequence of sgRNAs in this study
**Table S3:** Data corresponding to Figure 1C, 1D, and 1E
**Table S4:** Data corresponding to Figure 2B and 2C
**Table S5:** Data corresponding to Figure 4A and 4B
**Table S6:** Efficiency of plating assay, corresponding to Figure 5A
**Table S7:** Data corresponding to Figure 6A and 6C
**Table S8:** Data corresponding to Supplementary Figures
